# A Mechanism of Cohesin-Dependent Loop Extrusion Organizes Zygotic Genome Architecture

**DOI:** 10.1101/177766

**Authors:** Johanna Gassler, Hugo B. Brandão, Maxim Imakaev, Ilya M. Flyamer, Sabrina Ladstätter, Wendy A. Bickmore, Jan-Michael Peters, Leonid A. Mirny, Kikuë Tachibana-Konwalski

## Abstract

Fertilization triggers assembly of higher-order chromatin structure from a naïve genome to generate a totipotent embryo. Chromatin loops and domains are detected in mouse zygotes by single-nucleus Hi-C (snHi-C) but not bulk Hi-C. We resolve this discrepancy by investigating whether a mechanism of cohesin-dependent loop extrusion generates zygotic chromatin conformations. Using snHi-C of mouse knockout embryos, we demonstrate that the zygotic genome folds into loops and domains that depend on Scc1-cohesin and are regulated in size by Wapl. Remarkably, we discovered distinct effects on maternal and paternal chromatin loop sizes, likely reflecting loop extrusion dynamics and epigenetic reprogramming. Polymer simulations based on snHi-C are consistent with a model where cohesin locally compacts chromatin and thus restricts inter-chromosomal interactions by active loop extrusion, whose processivity is controlled by Wapl. Our simulations and experimental data provide evidence that cohesin-dependent loop extrusion organizes mammalian genomes over multiple scales from the one-cell embryo onwards.

**Highlights:**

- Zygotic genomes are organized into cohesin-dependent chromatin loops and TADs
- Loop extrusion leads to different loop strengths in maternal and paternal genomes
- Cohesin restricts inter-chromosomal interactions by altering chromosome surface area
- Loop extrusion organizes chromatin at multiple genomic scales

## INTRODUCTION

Chromatin is assembled and reprogrammed to totipotency in the one-cell zygote that has the potential to generate a new organism. The chromatin template upon which higher-order structure is built in the embryo is different for the maternal and paternal genomes at the time of fertilization. The maternal genome is inherited from the meiosis II egg in which chromosomes are condensed in a mitotic-like state (see **Figure 1A**). In contrast, the paternal genome is contributed from compacted sperm chromatin that is extensively remodeled as protamines are evicted and naïve nucleosomal chromatin is established (Rodman et al. 1981). The two genomes are reprogrammed as separate nuclei with distinct epigenetic signatures at the zygote stage (van der Heijden et al., 2005; Torres-Padilla et al, 2006; Mayer et al., 2000; Oswald et al., 2000; Ladstaetter and Tachibana-Konwalski, 2016). With the exception of imprinted loci, differences in chromatin states are presumably eventually equalized to facilitate the major zygotic genome activation (ZGA), which occurs at the 2-cell stage in mice (Aoki et al., 1997; Hamatani et al., 2004; Inoue et al., 2017). The establishment of zygotic genome architecture is therefore likely important for transcriptional onset and embryonic development.

**Figure 1:**
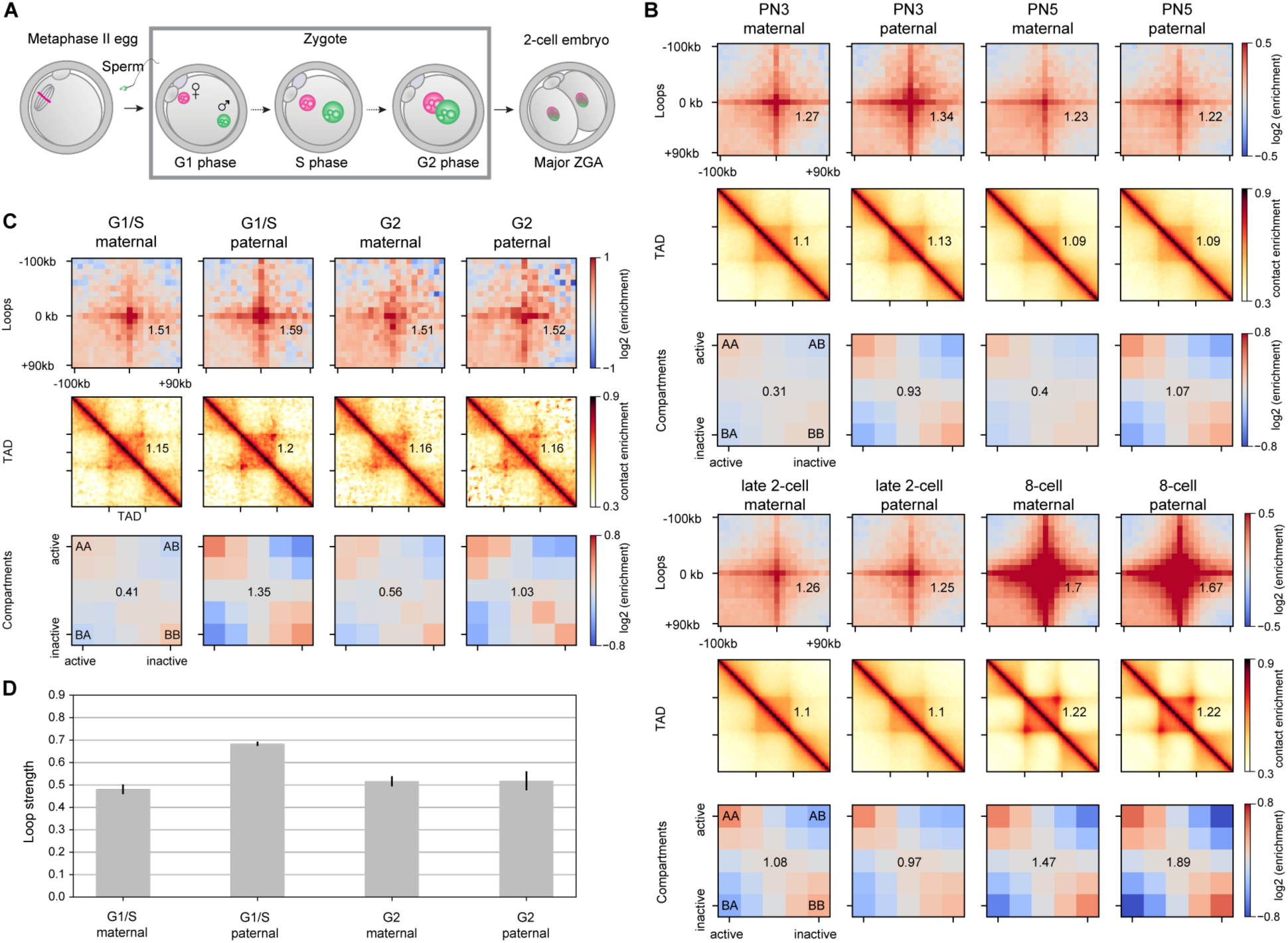
Zygotic chromatin is organized into loops, TADs and compartments that change during the first cell cycle. A) Embryonic development from fertilization of the metaphase II egg by sperm to zygote formation and division to the 2-cell embryo. Maternal and paternal genomes form separate nuclei in the zygote. The major zygotic genome activation (ZGA) occurs in the 2-cell mouse embryo. B) Average chromatin loops, TADs and compartments are detectable in maternal and paternal chromatin from the one-cell embryo onwards; data re-analyzed from Du et al., 2017. Zygotic pronuclear stage 3 (PN3) and stage 5 correspond to S and G2 phases, respectively. C) The strength of average loops, TADs and compartments becomes more similar between the maternal and paternal genomes as the zygotic cell cycle progresses (our data). D) Separation of of chromatin loops by size in G1/S and G2 phase maternal and paternal chromatin of zygotes. Loop strengths are the reported as the fractional enrichment above background levels (see Supporting Materials). Error bars displayed are the 95% confidence intervals obtained by bootstrapping.

Higher-order chromatin structures including: chromatin loops, topologically associating domains (TADs) and compartmentalization of active and inactive chromatin, are established during embryonic development (Flyamer et al., 2017; Hug et al., 2017; Du et al., 2017; Ke et al., 2017). Using single-nucleus Hi-C (snHi-C), we previously identified the presence of loops and TADs in mouse zygotes (Flyamer et al., 2017) by averaging contact maps over the positions of known TADs and loops (Rao et al., 2014). At variance with these findings, weak and “obscure” domain structures were detected by bulk Hi-C of early embryos. These results are difficult to interpret in the absence of comparable data with low cell numbers, such as in the interphase oocytes that are expected to have TADs and loops; thus, the absence of these structures may be due to technical reasons such as paucity of material. Interestingly, no TADs or loops are detected either in the rapidly dividing nuclei in early *Drosophila* embryos (Hug et al., 2017), where a substantial fraction of cells are in mitosis, or in metaphase II oocytes (Du et al., 2017; Ke et al., 2017). The lack of TADs and loops in mitotic chromosomes, however, is well-established from work in HeLa cells, suggesting no association to zygotic development (Naumova et al., 2013). Therefore, which higher-order chromatin structures are assembled in mammalian zygotes remains unresolved and the mechanisms that establish these structures in embryos are not known.

Studies in other cell types are beginning to provide insights into possible mechanisms leading to the establishment of higher-order chromatin structures. An early stepping stone towards understanding chromatin structure was the unexpected finding that the cohesin complex, known to be essential for sister chromatid cohesion, is expressed in post-mitotic cells (Wendt et al., 2008). Cohesin is a tripartite ring consisting of Scc1-Smc3-Smc1 that is loaded onto chromatin by the loading complex composed of Nipbl/Scc2 and Mau2/Scc4 and is released from chromosomes by Wapl (Ciosk et al., 2000; Tedeschi et al, 2013; Gandhi et al, 2006; Kueng et al, 2006). Mutations in Nipbl cause Cornelia de Lange Syndrome (CdLS) that is characterized by gene expression defects and altered chromatin compaction but no obvious defects in sister chromatid cohesion (Deardorff et al., 2007; Krantz et al., 2004; Musio et al., 2006; Tonkin et al., 2004; Nolen et al., 2013). Therefore, the idea emerged that cohesin may have roles beyond holding sister chromatids together. The discovery that cohesin colocalizes with CTCF and mediates its transcriptional insulation led to the conceptual advance that cohesin may hold DNA together not only between sister chromosomes but also in cis, within chromosomes (Wendt et al., 2008; Parelho et al., 2008). This is supported by the finding that depletion of Wapl leads to an increased residence time of chromosome-bound cohesin; moreover, it causes the formation of prophase-like chromosomes with cohesin-enriched axial structures termed “vermicelli” in G0 cells, and affects chromosome condensation (Tedeschi et al., 2013; Lopez-Serra et al., 2013). This discovery suggested that cohesin organizes intra-chromatid loops in interphase.

At the same time, chromosome conformation capture (3C)-based methods described interphase TAD structures with cohesin and CTCF enrichment at the boundaries (Dixon et al., 2012; Rao et al., 2014; Vietri Rudan et al., 2015). These observations led to the testable prediction that cohesin is required for TAD formation. However, cohesin depletion approaches including HRV protease-mediated cleavage, siRNA knockdown or conditional genetic knockout in cycling and differentiated cells only had minor effects on chromatin structure (Zuin et al.., 2013; Sofueva et al., 2013; Seitan et al., 2013), suggesting that cohesin was either not essential for TAD formation or protein depletion was inefficient. This caveat has recently been overcome by genetically knocking out the cohesin loading complex subunit Nipbl in post-mitotic liver cells as reported in a preprint (Schwarzer et al. 2016) and Mau2 in HAP1 cells (Haarhuis et al., 2017). However, cohesin was not directly manipulated in these studies, and it is assumed that cohesin loading is the sole function of Nipbl/Mau2. A recent preprint (Rao et al., 2017) tests the effect of auxin-inducible cohesin degradation on chromatin structure in a cancer cell line. The findings corroborate results showing reduced TAD and loop strengths after cohesin loading complex depletion obtained by Schwarzer et al., and Haarhuis et al.

A mechanism explaining the formation of TADs and loops is provided by the loop extrusion model. In this model (Fudenberg et al., 2016; Sanborn et al., 2015), dynamic chromatin loops are created in cis by loop extruding factors (LEFs). When a LEF binds to chromatin, it starts to translocate along the fiber in both directions, connecting successively further points, thus extruding a loop. Translocation of loop extruders is hindered by boundary elements often located at TAD boundaries. Loop extrusion naturally recapitulates the enrichments of contact probability in the corners of TADs, which are commonly referred to as “loops”, or “CTCF-mediated loops”. Cohesin is hypothesized to act as a loop extruder in interphase, while CTCF is likely the most prominent boundary element (Fudenberg et al., 2016; Sanborn et al., 2015; Nora et al., 2017).

Here we provide evidence that cohesin-dependent loop extrusion organizes higher-order chromatin structures of mammalian zygotic genomes. We show that cohesin is essential for formation of chromatin loops and TADs but not compartments and other large-scale zygote-specific structures in one-cell embryos. We find that inactivating cohesin release by Wapl depletion exacerbates differences in loop strengths between the maternal and paternal genomes that may be related to reprogramming. Remarkably, simulations indicate that most differences in global organization between the two zygotic genomes can be driven by changes in cohesin density and loop extrusion processivity. We further discovered that cohesin influences inter-chromosomal interactions by compacting chromatin; simulations indicate that this effect is due to altering the effective surface of chromosomes. We propose that cohesin-dependent loop extrusion organizes chromatin at multiple genomic scales from the mammalian one-cell embryo onwards.

## RESULTS

### Loops, TADs, and Compartments are Formed as Early as in One-Cell Embryos

We have previously shown that mouse zygotic genomes are organized into chromatin loops, TADs and compartments as early as G1 phase (Flyamer et al., 2017). Recently, it has been reported in bulk Hi-C of zygotes that few or “obscure” TAD structures are visible until around the 8-cell stage (Du et al., 2017; Ke et al., 2017). To resolve the confusion arising from these conflicting results, we re-analyzed these recent data (Du et al., 2017). Since loop and TAD locations are generally conserved across cell types (Rao et al., 2014; Dixon et al., 2012), but are unknown in zygotes, we used a list of known loop loci identified in CH-LX12 cells (Rao et al., 2014). For Hi-C data on low numbers of cells (Ulianov et al., 2017), TADs and loops are most visible when averaged over multiple positions (Flyamer et al., 2017). Contrary to the recent claims, our re-analysis of these data revealed that loops and TADs are present in 8-cell, 2-cell and even 1-cell embryos (**Figure 1B**). Thus, we affirm that zygotic genomes are folded into higher-order chromatin structures.

### Zygotic Genome Architecture Changes During the First Cell Cycle

Higher-order chromatin structure is established *de novo* for paternal chromatin and re-established after chromosome condensation for the maternal genome in zygotes. We noted that in G1 phase, loops differed in strength between genomes, with stronger loops visible in paternal chromatin (**Figures 1B-D**). It was conceivable that the differences observed in G1 are transient and that loops, TADs and compartments change during the first cell cycle. To test this, we performed snHi-C of nuclei isolated from G2 phase zygotes (**Figure 1C**). We found that zygotic genomes are organized into TADs, loops and compartments in G2 (**Figure 1C**), like in G1 phase. However, the average loop and TAD strengths had equalized between the genomes by G2 phase (**Figures 1B-D**). To investigate the source of different G1 loop strengths, we classified loops into small (100-150 kb), intermediate (150-250 kb) and large (250-500 kb), and computed average loops for each distance. We found that paternal chromatin has higher contact frequency primarily for small length loops in G1 (**Figure S1**), which could be a consequence of loop formation following protamine-histone exchange on sperm chromatin.

To determine whether compartmentalization also changes throughout the first cell cycle, we compared compartment strengths for our G1 and G2 phase data (**Figure 1C**). Consistent with the two recent reports (Du et al., 2017; Ke et al., 2017) (**Figure 1B**), the differences in compartmentalization of the maternal and paternal genome in G1/S phase zygotes were reduced by G2. However, maternal compartments were still weaker than paternal compartments. Likewise, we saw the initial differences in TAD strengths disappear throughout the cell cycle in two independent data sets (**Figure 1B** and **Figure 1C**). We thus conclude that initial differences in loop, TAD, and compartmentalization strengths between zygotic maternal and paternal genomes become less evident by the end of the first cell cycle.

### Cohesin is Essential for Zygotic Chromatin Folding into Loops and Domains

To gain insights into the mechanisms that generate zygotic genome architecture, we tested whether the candidate loop extruding factor cohesin is required for the formation of loops and domains. We used a genetic knockout approach based on *(Tg)Zp3*-Cre that deletes conditional alleles during the weeks of oocyte growth and generates maternal knockout zygotes after fertilization (**Figure 2A**). We have previously shown that Scc1 protein is efficiently depleted and sister chromatid cohesion fails to be established in *Scc1*^Δ*(m)/+(p)*^ zygotes (hereafter referred to as *Scc1*^Δ^ according to the maternal allele) (**Figure S2A**, Ladstätter & Tachibana-Konwalski, 2016). Since sister chromatid cohesion is maintained by Rec8-cohesin in oocytes (Tachibana-Konwalski et al., 2010; Burkhardt et al., 2016), Scc1 depletion has no effect on chromosome segregation prior to fertilization and therefore a clean Scc1-cohesin knockout zygote is generated.

**Figure 2:**
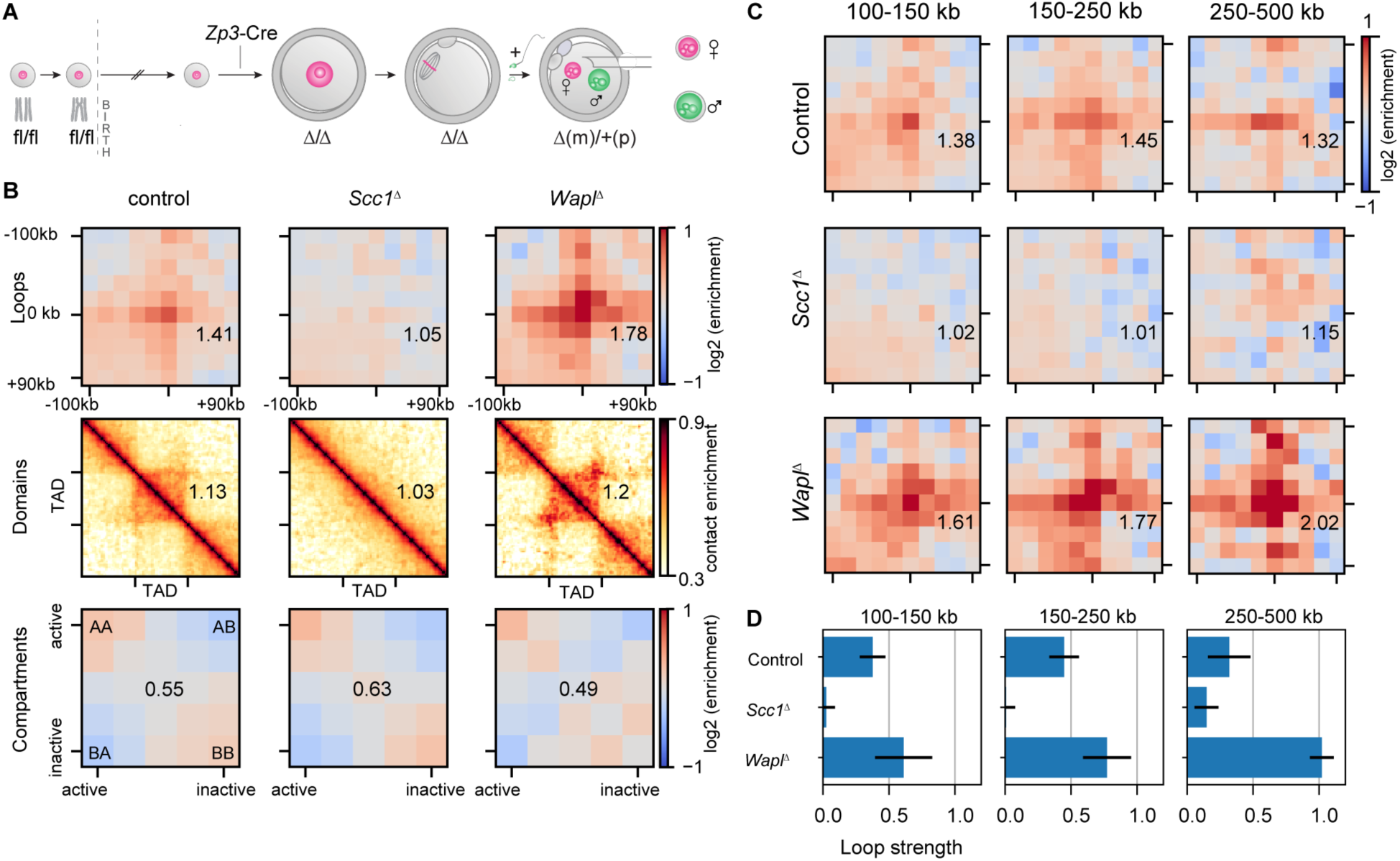
Conditional genetic knockouts of *Scc1* and *Wapl* reveal cohesin’s essential role in formation of loops and TADs in mouse zygotes. A) Generation of conditional genetic knockout oocytes by *Zp3*-Cre recombinase in post-recombination growing phase mouse oocytes. Fertilization produces maternal knockout zygotes (maternal, m, paternal, p, alleles). Maternal and paternal nuclei are extracted from zygotes before being subjected separately to snHi-C. B) Average loops, TADs and compartments in control (*Wapl*^*fl*^ *and Scc1*^*fl*^), *Scc1*^*Δ*^ and *Wapl*^*Δ*^ zygotes. Both maternal and paternal data are shown pooled together. C) Separation of loops by size for control, *Scc1*^*Δ*^ and *Wapl*^*Δ*^ zygotes for maternal and paternal data pooled together. D) Loop strengths are reported as the fractional enrichment above background levels (see Supporting Materials). Error bars displayed are the 95% confidence intervals obtained by bootstrapping pooled single cell loops.

To test whether Scc1 is essential for the formation of TADs and loops in zygotes, we performed snHi-C (Flyamer *et al*, 2017) on genetically modified embryos. Both chromatin structures were detectable in control *Scc1*^*fl*^ zygotes (**Figure 2B**). Remarkably, TADs and loops were largely, if not entirely, absent in *Scc1*^Δ^ zygotes, both in maternal and paternal nuclei (**Figure 2B**; **Figure S2B**). Compartmentalization of active and inactive chromatin from both maternal and paternal genomes was increased over 2-fold in *Scc1*^Δ^ compared to controls (**Figure S2B**). We conclude that cohesin is essential for loop and domain formation and antagonizes compartmentalisation, which is consistent with the notion that independent and possibly competing mechanisms generate these higher-order chromatin structures (Nora et al., 2017; Schwarzer et al., 2016; Haarhuis et al., 2017).

### Wapl Controls the Size of Cohesin-Dependent Chromatin Loops

The loss of loops and domains in the absence of cohesin could either be due to an indirect effect, for example on gene expression, or reflect a direct requirement for cohesin’s involvement in loop formation. The loop extrusion model predicts that increasing the residence time of cohesin on chromosomes strengthens existing loops and promotes the formation of longer loops in a population of cells (Fudenberg et al., 2016). The residence time of cohesin on chromatin can be increased by more than ten-fold by inactivating cohesin release through Wapl depletion (Tedeschi et al., 2013). To test whether loop and domain structures in zygotes are enhanced by inactivating release of chromosomal cohesin, we generated *Wapl*^*Δ(m)/+(p)*^ *(Wapl*^*Δ*^) zygotes using the same strategy as described for *Scc1* (**Figure 2A**). We performed snHi-C of S/G2 phase *Wapl*^*Δ*^ zygotes and compared these to pooled control data from *Wapl*^*fl*^ and *Scc1*^*fl*^ zygotes, which are wild-type for *Wapl*. Both loops and domains are stronger in *Wapl*^*Δ*^ compared to control zygotes (**Figure 2B; Figure S2C**), in agreement with what has been observed in *Wapl*^*Δ*^ HAP1 cells (Haarhuis et al. 2017). We conclude that cohesin release from chromosomes by Wapl is essential for regulating loop and domain formation.

We next tested whether inactivating cohesin release from chromosomes causes changes to average loop strengths, and increased processivity of loop extruders. We found that loops are stronger in pooled *Wapl*^Δ^ zygote data compared to controls for all tested genomic distances (**Figure 2C and 2D**). Interestingly, unlike for controls, where loop strength was invariant with increasing distance, in *Wapl*^Δ^ zygotes they increased from short to large distances with up to 100% enrichment of contacts above background levels (**Figure 2D**). These results are consistent with the loop extrusion model and suggest that in wild-type cells, Wapl limits the extent of loop formation by releasing cohesin from chromosomes, impeding its processivity. Altogether, we conclude that cohesin directly regulates loop and domain formation in the one-cell embryo.

### Cohesin organizes chromosomes at the sub-Megabase scale

To further investigate the mechanism by which cohesin shapes genome architecture, we studied the genome-wide contact probability, P_c_(s), for chromatin loci separated by genomic distances, s. Consistent with our previous observations of wild-type zygotes (Flyamer et al., 2017), control cells have a P_c_(s) curve that changes slowly below 1 Mb, reflecting local chromatin compaction; it changes steeply at or after 1 Mb in paternal chromatin and exhibits another plateau near 10 Mb in maternal chromatin, likely reflecting long-range chromatin interactions remaining from compaction to the mitotic state (**Figure 3A**) (Flyamer et al., 2017; Naumova et al., 2013). Interestingly, the P_c_(s) curve of *Scc1*^*Δ*^ zygotes lost the shallow <1 Mb region, and followed a power law of s^-1^.^5^, up to 1 Mb in both maternal and paternal genomes; the power law stretched up to 10 Mb in paternal chromatin (**Figure 3B**). This indicates that in the absence of cohesin, zygotic chromatin resembles a three-dimensional random walk as previously observed in yeast (Tjong et al., 2012; Mizuguchi et al., 2014; Halverson et al., 2014). Conversely, in *Wapl*^*Δ*^ zygotes, the contact probability was enriched and more shallow up to ∼300 kb further than in controls (**Figure 3C**). In both *Scc1*^*Δ*^ and *Wapl*^*Δ*^ P_c_(s) curves, contact probability features at >10 Mb remain largely unchanged. Therefore, differences in long-range interactions (>10 Mb) between maternal and paternal chromatin are cohesin-independent. Thus, we conclude that cohesin is directly involved in shaping the P_c_(s) curve up to ∼1 Mb, and its effect is a deviation in contact probability above the s^-1^.^5^ power-law in mouse zygotic chromatin.

**Figure 3:**
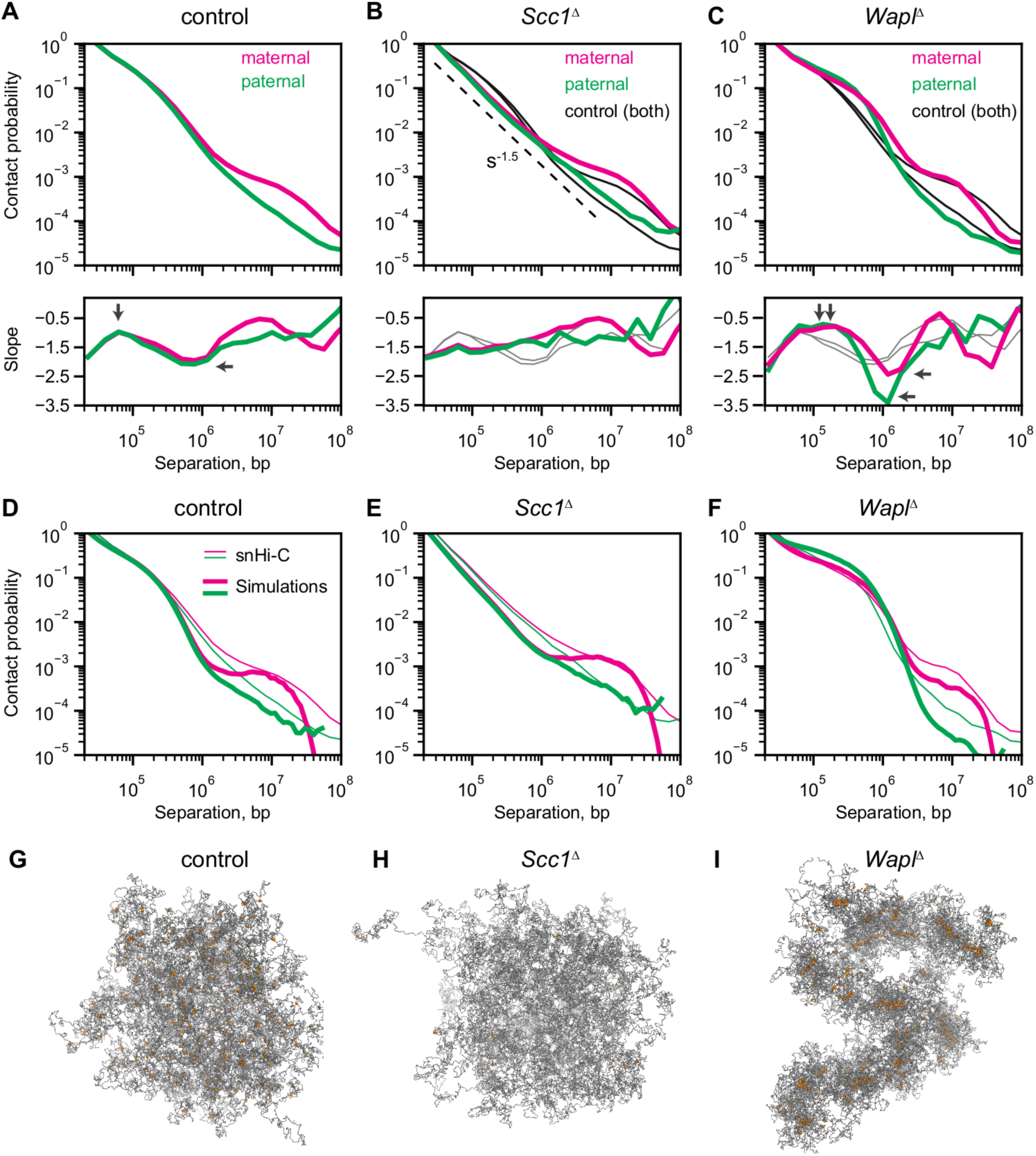
Differences in P_c_(s) between conditions. A-C) Experimental P_c_(s) for maternal and paternal chromatin for control, *Scc1*^*Δ*^, and *Wapl*^*Δ*^ conditions. Black solid lines in B and C show the control curves as a reference to guide the eye. Slopes of the log(P_c_(s)) curves for each condition are are shown in the sub-panel below each P_c_(s) plot. Vertical arrows on the slope subpanels indicates the inferred average size of cohesin extruded loops. Horizontal arrows on the slope panels indicate whether there are expected differences in cohesin linear density on the chromatin; whereas in controls, C, little difference in cohesin density is expected. D-F) Simulated chromatin P_c_(s) for the control, *Scc1*^*Δ*^, and *Wapl*^*Δ*^ conditions. Simulation P_c_(s) curves shown in thick lines and experimental P_c_(s) curves in thin lines. G-I) Representative images of the simulated paternal chromatin fiber used for the P_c_(s) calculations in Panels D-E. The chromatin fiber is coloured in gray, and the locations of the cohesins coloured in orange.

To help elucidate the mechanism of loop formation by cohesin, we developed a new method for analysis of P_c_(s) curves aiming to derive sizes of extruded loops and density of cohesin molecules from these curves. We developed and tested this method using polymer simulations of loop extrusion, where sizes of loops and cohesin density are either set or can be directly measured. Our analysis suggests that it is possible to estimate both the average length of loops created by cohesins, and cohesin density by studying the derivative of the P_c_(s) curve in log-log space (**Figure S3A**): The location of the maximum of the derivative curve (i.e. position of the smallest slope) closely matches the average length of extruded loops, and the depth of the local minimum at higher values of *s* increases with the number of loop-extruding cohesin in simulated chromatin (**Figure S3A**). Note that sizes of extruded loops are smaller than the processivity of each cohesin, defined as the loop size extruded by unobstructed cohesin, suggesting some degree of crowding of cohesins on DNA, as expected theoretically (Fudenberg et al., 2016; Goloborodko et al.,2016). To validate this approach, we performed this analysis on recent experimental data of *Wapl*^*Δ*^ and control HAP1 cells (Haarhuis et al., 2017); we report good agreement between our predictions for loop size and relative cohesin numbers by the log(P_c_(s)) slope analysis by matching simulation results to experimental data (**Figure S3B**).

Interpreting our zygote data in light of the log(P_c_(s)) slope analysis, we estimated that loop extrusion by cohesin results in the average loop length of 60-70 kb in controls. In contrast, in *Wapl*^*Δ*^ zygotes, the length of loops extruded by cohesin was doubled to 120 kb. Indeed, varying cohesin density and processivity parameters in our simulations and generating P_c_(s) curves, we find that experimental P_c_(s) curves are best matched by these values (**Figures 3D-F**): We obtain a 74 kb average loop length for the model of control zygotes, and 111 and 165 kb for *Wapl*^*Δ*^ for paternal/maternal models. Corresponding, respectively, with a processivity of 120 kb in control and 480 kb in *Wapl*^*Δ*^ conditions, a mean cohesin separation of 120 kb in control and maternal *Wapl*^*Δ*^, and 60 kb in paternal *Wapl*^*Δ*^ conditions. We also made similar estimates for population Hi-C for *Wapl*^*Δ*^ and control HAP1 cells (Haarhuis et al., 2017), showing good agreement with results for zygotes. An apparent discrepancy arises between the predicted average length of an extruded loop (<150 kb) versus a loop visible on the Hi-C map (seen as far as 1 Mb). We can make sense of this by noting that the Hi-C loops visible at 1 Mb are likely formed of a collection of much shorter cohesin extruded loops that have bumped into each other. However, because the extrusion process is stochastic, the location where the cohesin extruded loops bump into each other will be variable, so we do not necessarily see “Hi-C loops” at the locations where they stall. Thus, this new method for analysis of P_c_(s) curves provides a framework for the interpretation of genome-wide contact probability in mammalian cells, and we conclude that Wapl is regulating cohesin processivity.

### Loop extrusion leads to differences in compaction of maternal and paternal chromatin

We next considered whether loop extrusion differs between the maternal and paternal genomes. Extending our analysis of contact probabilities to study differences in maternal and paternal genomes, we find that the sizes of extruded loops and number of cohesins are similar in controls. In contrast, in *Wapl*^*Δ*^ zygotes, the log(P_c_(s)) derivative analysis (**Figure 3C**) and our simulations (**Figure 3F**) indicate that paternal chromatin may have the same processivity of cohesins but higher numbers of cohesins, albeit resulting in loops that are shorter, in part due to crowding. Insulation scores (see **Supporting Materials**) provide a rationale for shorter loops in the paternal genome as they suggest that loop borders are stronger boundary elements for cohesin in paternal chromatin (**Figure S2D**). Likewise, quantifying the loop strength above background separated by distance also suggests that paternal loops are stronger than maternal loops at short distances upon Wapl depletion (**Figure S2B**). This difference in abundance of chromatin-associated cohesin between maternal and paternal genomes is revealed when cohesin unloading is abolished, suggesting differences in cohesin loading rates between maternal and paternal chromatin. Stronger enrichment of interactions in Hi-C loops, higher insulation, and possibly higher cohesin loading rates may all reflect the transcriptionally active state specific for paternal chromatin (Adenot et al., 1997), and indicate that transcription is not required for loop extrusion in the maternal genome.

We next tested whether differences in loops between maternal and paternal chromatin lead to changes in chromatin compaction by microscopy. We expressed Scc1-EGFP in *Scc1*^*Δ/Δ*^*Wapl*^*Δ/Δ*^ oocytes to ensure that all Scc1 present in the cell is fluorescently tagged and thus increase the sensitivity of vermicelli detection (**Figure 4A**). Meiosis II oocytes were fertilized *in vitro* and zygotes were imaged by time-lapse microscopy. Scc1-EGFP accumulated in both nuclei at roughly the same time and was uniformly diffuse in the nuclei of wild-type zygotes (**Figure 4B and 4C; Supplementary Movie 1; Supplementary Figure S4A and S4B**). In contrast, Scc1-EGFP vermicelli were detected in both nuclei of *Scc1*^Δ^*Wapl*^Δ^ zygotes, with more prominent, brighter structures appearing in the maternal nucleus (3/3 zygotes) (**Figure 4B and 4C; Supplementary Figure S4C; Supplementary Movie 2**). The formation of vermicelli occurs prior to the major ZGA (Aoki et al., 1997; Hamatani et al., 2004), consistent with the idea that transcription is not required for loop formation. We conclude that inactivation of cohesin release leads to formation of vermicelli that are morphologically distinct between maternal and paternal chromatin.

**Figure 4:**
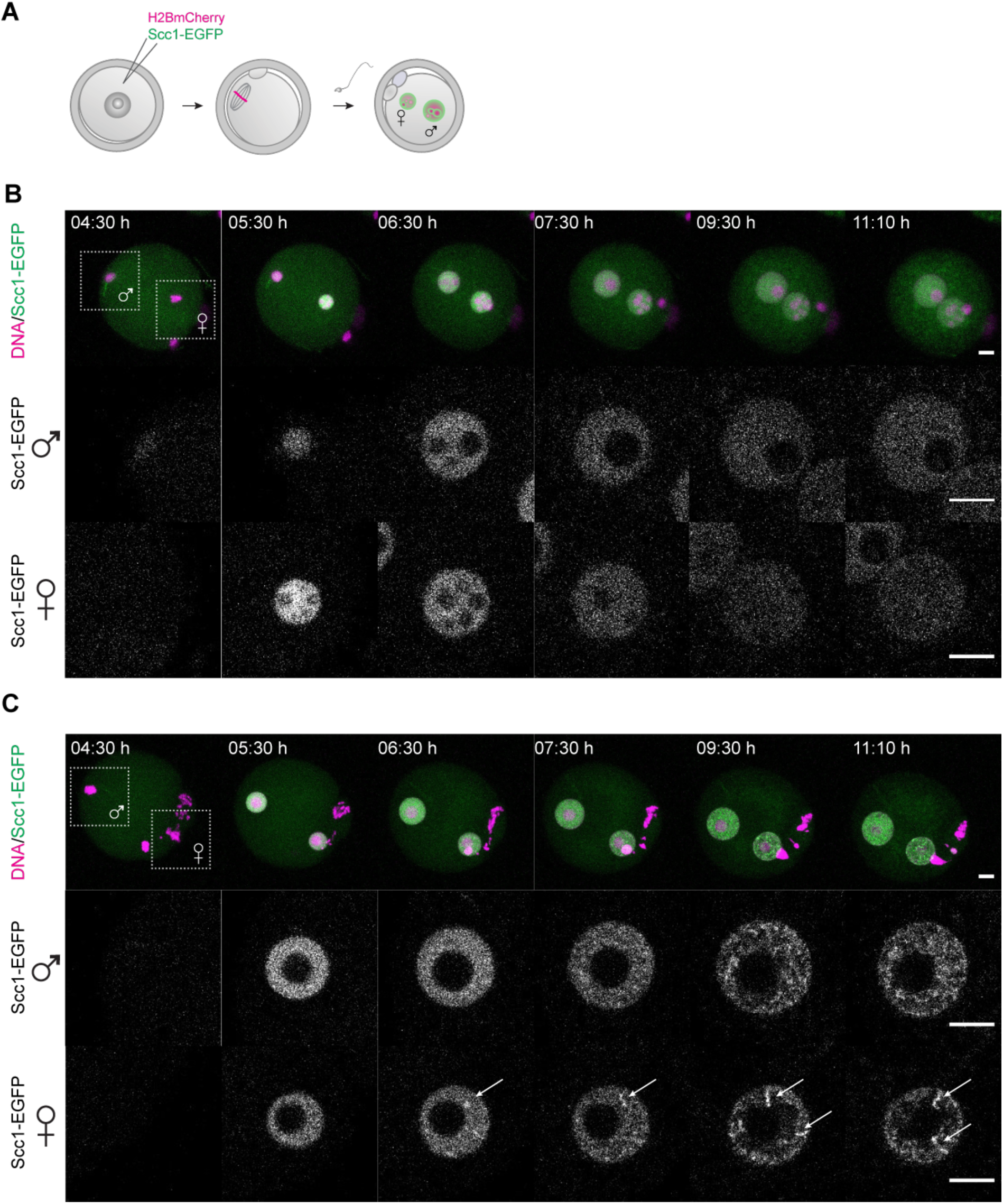
Live-cell imaging of vermicelli formation in wildtype and *Scc1ΔWaplΔ*zygotes expressing Scc1-EGFP and H2B-mCherry. A) Germinal vesicle-stage oocytes were injected with mRNA encoding H2B-mCherry to mark chromosomes (magenta) and Scc1-EGFP to label vermicelli (green), matured to meiosis II, fertilized *in vitro* and followed by time-lapse microscopy. B) Still images of live wild-type zygotes expressing Scc1-EGFP and H2B-mCherry. Above: Z-stack maximum intensity projection of zygotes. Below: Z-slices of the cropped areas showing paternal and maternal nuclei separately. Images were adjusted in brightness/contrast in individual imaging channels in the same manner for (B) and (C). Scale bar: 10 μm. C) Still images of live *Scc1*^*Δ*^*Wapl*^*Δ*^ zygotes expressing Scc1-EGFP and H2B-mCherry. Above: Z-stack maximum intensity projection of zygotes. Below: Z-slices of the cropped areas showing paternal and maternal nuclei separately. Arrows indicate Scc1-EGFP enriched structures. Scale bar: 10 μm.

We examined DNA morphology at higher resolution in fixed zygotes to quantify maternal and paternal chromatin compaction. Both maternal and paternal chromatin is compacted into vermicelli-like structures revealed most clearly in individual z-sections of *Wapl*^Δ^ zygotes, with a significant change in the coefficient of variation in intensity between control and *Wapl*^Δ^ zygotes (**Figure 5A and 5B, Figure S5A,** p-value=1.88*10^-7^). Strikingly, additional DAPI-intense structures were visible specifically in the maternal nucleus (n=19/23 zygotes), indicating a higher degree of compaction in maternal than paternal chromatin. These structures likely correspond to the vermicelli observed in time-lapse movies (**Figure 4B and 4C; Supplementary Figure S4C; Supplementary Movie 2**). Quantification of the texture in images using the grey-level co-occurrence matrices revealed that the contrast between pixels is stronger in maternal than paternal nuclei (**Figure 5C and Figure S5B-D**), implying a more structured and less homogeneous nuclear architecture. We cannot distinguish whether this reflects solely the “brighter” structures specific to maternal chromosomes or a quantitative genome-wide difference in vermicelli between the two nuclei. We suggest that inactivating cohesin release has a differential effect on chromatin compaction of maternal and paternal chromatin.

**Figure 5:**
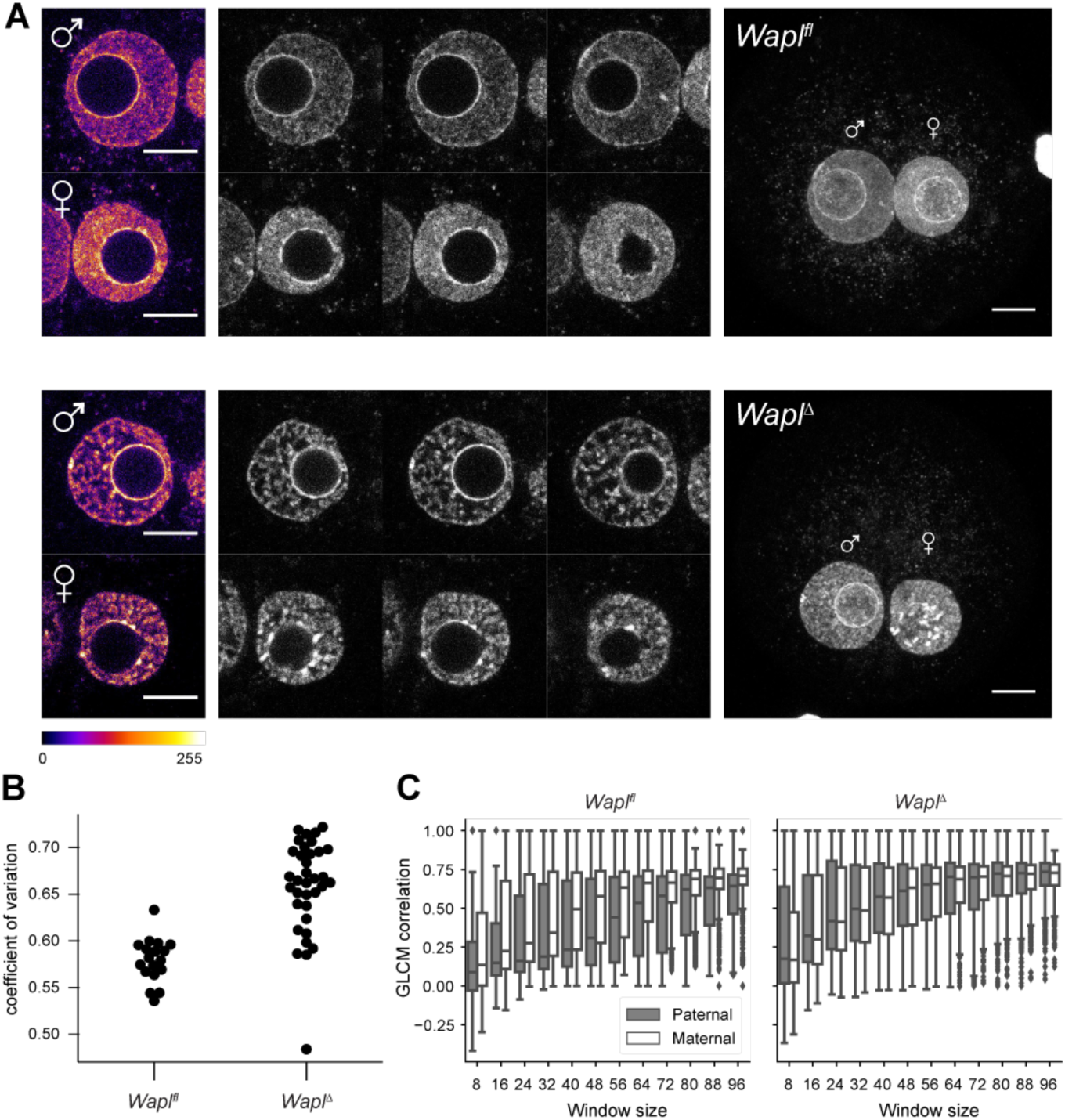
Distinct maternal and paternal chromatin compaction in *WaplΔ*zygotes. A) Representative images of paternal and maternal nuclei stained with DAPI of *Wapl*^*fl*^ (n=10) and *Wapl*^*Δ*^ (n=23) zygotes (see also Supplementary Figure 5). Top: *Wapl*^*fl*^, Bottom: *Wapl*^*Δ*^. Left: Cropped z-slices from the middle section of the nucleus in fire look up table (LUT). Middle: Cropped z-slices of nuclei separated by 3 μm. Right: Maximum Intensity Projection (MIP) of zygotes. Images were adjusted in brightness/contrast settings using ImageJ. Settings were adjusted for z-slices and MIP individually but in the same manner for *Wapl*^*fl*^ and *Wapl*^*Δ*^ zygotes. Scale bar: 10 μm. B) Coefficient of variation of DAPI intensity for nuclei of *Wapl*^*fl*^ (n=18) and *Wapl*^*Δ*^ (n=35) zygotes (p-value=1.88*10^-7^). C) Boxplots showing GLCM contrast (local variation of intensity) in paternal (grey) and maternal (white) nuclei in *Wapl*^*fl*^ (n=10) and *Wapl*^*Δ*^ (n=12) zygotes with increasing window sizes. All images were adjusted in brightness/contrast in the individual imaging channels using ImageJ. Cropped areas are indicated. Scale bar: 10μm.

To corroborate the findings obtained by microscopy, we used the polymer simulations that best matched experimental scalings to test whether the 3D organization of cohesins displayed preferentially “axially enriched” structures. We found consistently that vermicelli are more visible in the paternal simulations; at odds with expectations, maternal chromatin formed weaker vermicelli (**Figure S3C**). This result suggests that some other elements beyond our simple model of loop extrusion may be at play for the formation of vermicelli in live cells. Nevertheless, both our snHi-C data and microscopy show that loop formation differs for zygotic maternal and paternal genomes when cohesin release is prevented by Wapl depletion. By regulating cohesin release, Wapl thus maintains interphase chromatin in a less compact state; moreover, it helps restrict the size of extruded cohesin loops to shorter length scales.

### Effect of cohesin and loop extrusion on global chromosome organization

Population and single-cell Hi-C studies have revealed that interactions between non-sister chromatids are diminished during mitosis (Naumova et al., 2013; Nagano et al., 2017). A possible interpretation is that a more compact, linearly ordered chromosome directly affects the frequency of inter-chromosomal interactions. To investigate whether the vermicelli chromosomes are more mitotic-like, and to test whether cohesin might play a role in chromosome compaction, we quantified the levels of inter-chromosomal contacts, hereby called trans-contacts, in zygotic chromatin by snHi-C (**Figure 6A, see Supporting Methods**). We found inter-chromosomal contact frequencies of 8% for both nuclei in interphase (G1/S or G2), which are consistent with values reported for mouse ES cells at a similar cell cycle stage (Nagano et al., 2017). Interestingly, *Wapl*^*Δ*^ zygotes had reduced trans interaction fractions, with a mean value of 6%, closer to values reported for early G1 (Nagano et al., 2017), but were not significantly different than controls (p<0.2, Mann-Whitney U-test). In contrast, *Scc1*^*Δ*^ cells had significantly larger trans interaction fractions as compared to controls (**Figure 6A**; an over 40% increase, p<0.02 Mann-Whitney U-test). These results suggest a novel role for chromosomal Scc1-cohesin in reducing interaction frequencies between non-sister chromatids.

**Figure 6:**
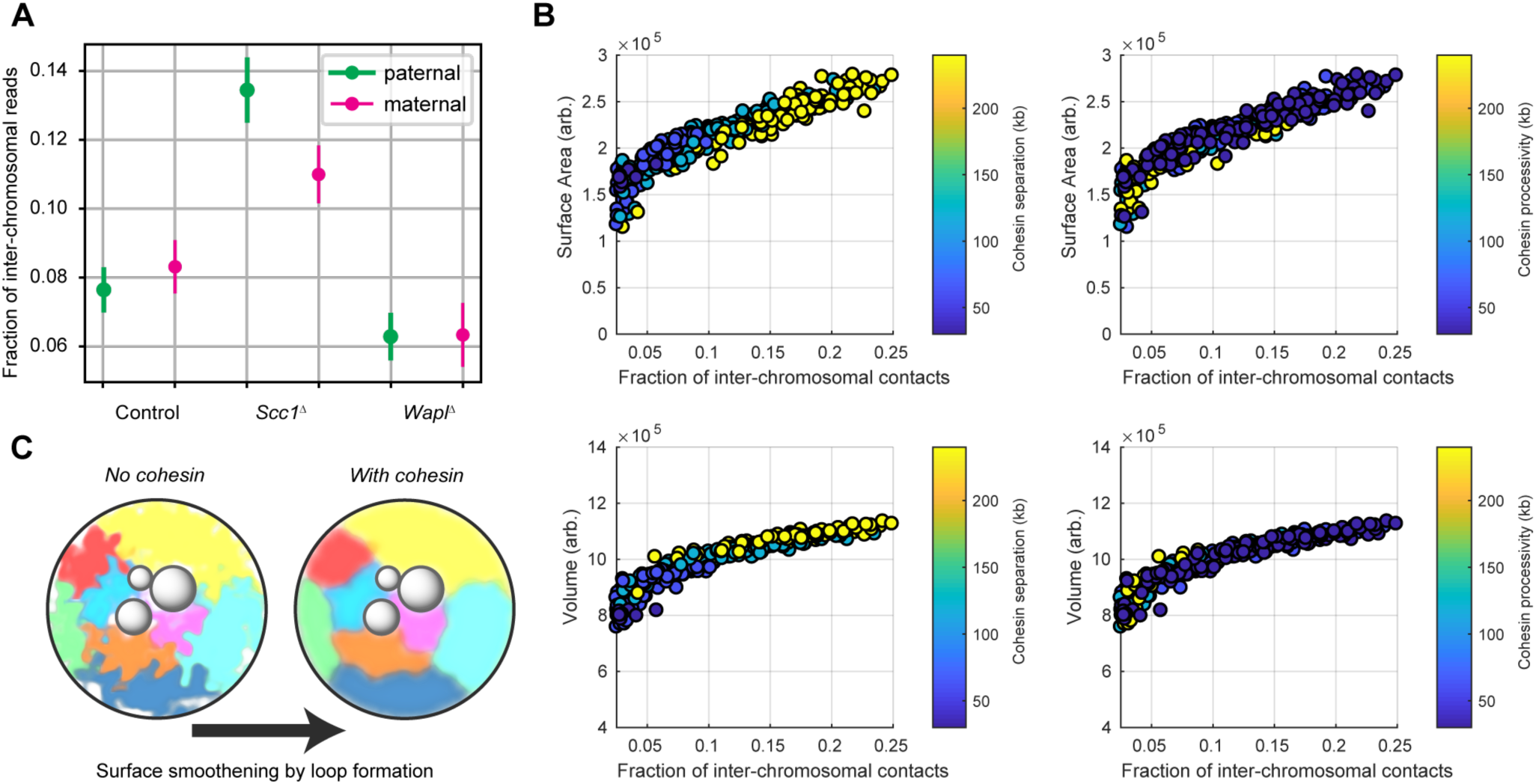
The influence of cohesin on inter-chromosomal contacts. A) The number of snHi-C contacts mapping to regions on distinct chromosomes, as a fraction of the total number of mapped contacts is shown for each of the experimental conditions. Error bars are the standard error of the mean. B) Spatial, geometric properties of simulated chromatin undergoing loop extrusion for different loop extrusion parameters. The fraction of inter-chromosomal contacts were calculated using a Hi-C cutoff radius of 5 monomers (75 nm). The surface area and volume of the simulated chromatin fiber were calculated from the concave hull, and used an effective radius for each monomer equal to the Hi-C cutoff radius (see **Supplemental Methods**). C) A schematic model illustrating that cohesin loop extrusion can modulate the surface area smoothness of chromosomes and reduce the frequency of inter-chromosomal interactions.

To investigate the mechanism by which cohesin modulates inter-chromosomal interactions, we turned to our polymer simulations of loop extrusion. We varied simulation parameters for processivity and linear density of cohesins and probed for changes in absolute numbers of contacts within and between chromosomes. An increase in density or processivity of cohesins resulted in an increase in intra-chromosomal contacts and a decrease in the absolute and relative counts of trans-chromosomal contacts (data not shown), consistent with experimental results. Thus, simulations suggest that cohesin can regulate the relative abundance of contacts between chromosomes by forming chromatin loops.

To better understand how loop formation that operates at the sub-megabase scale can affect inter-chromosomal contacts, we examined the effects of loop extrusion on the sizes of chromosomes and shapes of their surfaces. First, we tested whether more cohesin and loop extrusion can lead to greater chromatin compaction, a smaller chromosome volume, and therefore fewer interactions between neighbouring chromosomes. Chromosome size correlated with abundance of trans contacts, showing up to 30% change over the ranges expected for *Scc1*^*Δ*^ and *Wapl*^*Δ*^ (**Figure 6B**, **Figure S6B**). Interestingly, however, surface area, defined from the concave hull of modeled chromosomes (see Supporting Information), turned out to be more affected by loop extrusion and showed over 80% change over the range of expected values for our experiments (**Figure 6B**). Moreover, chromosome surface area is a good predictor of abundance of both inter- and intra-chromosomal contacts. By visualizing sample polymer conformations for low and high cohesin densities, we found that a decrease in the number of extruded loops leads to a surface roughening, whereas increased compaction by loop extrusion smoothens out the polymer surface. The density of extruded loops in simulated polymer models of chromosomes correlates better with the chromosomes’ surface area than the processivity of cohesin with surface area (**Figure 6B**). Based on these simulations, we propose a model whereby intra-chromosomal loops modulate chromatin surface area in interphase, which results in reduced contact frequencies between chromosomes (**Figure 6C**). Loop extrusion by cohesin therefore fosters intra-chromosomal interactions by creating cis loops and limits inter-chromosomal interactions by modulating surface area.

## DISCUSSION

We settle a decade’s long open question by demonstrating that cohesin is directly involved in forming chromatin loops and TADs. Cohesin was identified over two decades ago for its role in chromosome segregation, sister chromatid cohesion, and DNA damage repair (Peters et al., 2008). More recent studies have shown that cohesin colocalizes with CTCF and is associated with TADs and chromatin loops (Wendt et al., 2008; Dixon et al., 2012; Nora et al., 2012; Phillips-Cremins et al., 2013; Rao et al., 2014), which implicated cohesin as a regulator of intra-chromosomal structure. Since chromatin loops and TADs have functional roles in gene regulation, such as preventing aberrant expression of genes (Lupianez et al., 2015; Franke et al., 2016; Flavahan et al., 2015), it has become a major endeavour to understand to what degree cohesin is involved in shaping chromatin structure. Early studies directly degrading or knocking out cohesin showed only mild effects on chromatin structure (Seitan et al., 2013; Sofueva et al., 2013; Zuin et al., 2014).

We show that genetic deletion of the Scc1 subunit of cohesin in mouse oocytes abolishes formation of loops and TADs in the one-cell embryo. In contrast, chromatin loops are larger on average when cohesin release from chromosomes is prevented by Wapl depletion. Together, these results demonstrate that cohesin is essential for the formation of loops and TADs, and show that cohesin directly regulates their structure, consistent with two recent reports in human cell line studies. A recent preprint (Rao et al., 2017) shows loss of loops and TADs by acute degradation of Scc1/Rad21. Another study (Haarhuis et al., 2017) demonstrates that Wapl deletion leads to formation of longer loops. Similarly, a preprint reports that deletion of Nipbl led to disappearance of loops and TADs in post-mitotic liver cells (Schwarzer et al. 2016). We extend these insights into the fundamental principles of chromatin organization by providing a quantitative framework to understand the experimental results via polymer simulations. Our work also diverges from these reports in that we show cohesin is essential for forming loops and TADs starting from the one-cell embryo, which was hitherto unclear.

Crucially, our system enabled us to study how cohesin differentially affects the establishment of higher-order structure in maternal and paternal genomes that undergo reprogramming to totipotency. Interestingly, differences in maternal and paternal chromatin loops became more evident in *Wapl*^*Δ*^ zygotes. As in controls, paternal chromatin loops were stronger, and TADs more insulating than in maternal chromatin. Unlike controls, loop sizes differed by an estimated 60 kb, with longer loops present in the maternal genome. By microscopy, we also observed differences in global chromatin compaction between maternal and paternal genomes. We speculate that the differences are due to a combination of distinct epigenetic modifications and loop extrusion dynamics.

Our data strongly supports a model that cohesin forms loops and TADs by the mechanism of active loop extrusion (Fudenberg et al., 2016; Sanborn et al., 2015), and provides a quantitative rationale for the longer loop lengths in the *Wapl*^*Δ*^ zygotes. Our polymer simulations suggest that the key determinants for global genome organization by cohesin are their density and processivity, which is the product of residence time and extrusion velocity. Longer chromatin loop sizes in *Wapl*^*Δ*^ zygotes are quantitatively consistent with about a two-fold increase in cohesin processivity in the absence of Wapl, which results in about a 50% increase in the sizes of extruded loops as estimated from the derivative of log(P_c_(s)). Our present data does not distinguish whether increase in processivity reflects an increase in loop extruding speed, residence-time or both, but this is an interesting avenue for future research. Interestingly, sizes of extruded loops are smaller than processivity since extrusion is obstructed by interactions of boundary elements (with CTCF among them) and other chromatin-associated cohesins. In support of the model of active loop extrusion, Wang and coworkers recently provided the first direct *in vivo* evidence that condensins, which are related to cohesins, actively translocate on bacterial chromatin and align flanking chromosomal DNA (Wang et. al, 2017; Tran et al., 2017). A recent preprint has since demonstrated that eukaryotic yeast condensins are mechanochemical motors that translocate along DNA in an ATP-dependent fashion (Terekawa et al., 2017). Thus, it is likely that eukaryotic cohesins employ active loop extrusion to form chromatin loops and TADs, but we cannot rule out the possibility that accessory factors aid the extrusion process.

Two recent reports have provided conflicting views of the higher-order chromatin organization in mammalian embryos, suggesting that the mammalian zygote is largely unstructured (Du et al., 2017; Ke et al., 2017). In both studies, no or “obscure” TADs were detected in embryos before 8-cell stage (Du et al., 2017; Ke et al., 2017), where TAD were detected using insulation score and directionality index analysis (Dixon et al., 2012; Giorgetti et al., 2016) with a large window size (0.5-1 Mb). We note that non-zero insulation scores or directionality indices do not necessarily reflect the existence of a TAD since these metrics cannot distinguish TADs from compartments without other information; weak compartments in zygotes can further affect insulation scores or directionality indices. To settle whether or not TADs and loops exist in zygotes, we reanalyzed data from these studies. We demonstrated that TADs and loops can be clearly identified at all embryonic development stages, when known positions of TADs and loops were used; we also show that these structures depend critically on cohesin for their establishment.

There are several possible explanations for why loops and TADs are weaker in zygotes than for later stage embryos, such as weaker boundary elements, lower rate of cohesin loading, or lower cohesin processivity. The difference in processivity is unlikely as our analysis suggests similar processivity in paternal zygotes and HAP1 cells. Transcription, however, can affect loading and positioning of cohesin (Kagey et al., 2010; Busslinger et al., 2017). Since the paternal genome was shown to undergo transcriptional activation earlier than the maternal genome (Bouniol, 1995), this hypothesis predicts that loops may be shorter in paternal genomes, and TADs boundaries/loop strengths are stronger. In support of this idea, we show both TAD and loops strengths are greater in the paternal early G1 zygotes, but these differences disappear in G2 as both genomes approach the major ZGA. Interestingly, however, we find that the average length of loops extruded by cohesins is similar between both maternal and paternal genomes. We thus suggest that the weaker structural features seen in the zygotic genome arise due to either paucity of boundary elements for cohesin loop extrusion or increasing amounts of chromatin-associated cohesin.

Unexpectedly, we discovered that cohesin-dependent chromosome compaction reduces inter-chromosomal interactions in interphase. We therefore propose a model in which the surface roughness of chromosomes affects inter-chromosomal interactions and absence of cohesin leads to more interdigitation between chromosomes. We speculate as to what might be the functional consequences of increased inter-chromosomal interactions due to interphase chromosome decompaction. Given that topoisomerases cannot distinguish between DNA strands in cis and in trans, it is conceivable that increased number of trans interactions could lead to catenations that can be damaging during chromosome segregation. We therefore propose that the ancestral role of cohesin in forming intra-chromosomal loops during interphase could help promote proper chromosome segregation during cell division.

Our model of cohesin as a chromatin surface area regulator also raises important new points. If the active formation of loops can reduce inter-chromosomal interactions, then it is conceivable that loop formation creates local neighbourhoods on the chromatin fiber that also reduce the frequency of interactions with more distal segments of chromatin on the same chromosome. We speculate that the formation of loops can have important implications for reducing spurious enhancer-promoter looping interactions by reducing interdigitation between distant regions of the same chromosome.

In all, our work establishes which higher-order chromatin structures are built shortly after fertilization in the mammalian zygote. The differences in maternal and paternal loops generated by cohesin-dependent loop extrusion provide an entry point to understanding how the two genomes change from a transcriptionally silent and terminally differentiated state to a transcriptionally active and totipotent embryonic state.

## SUPPLEMENTARY FIGURES

**Figure S1:**
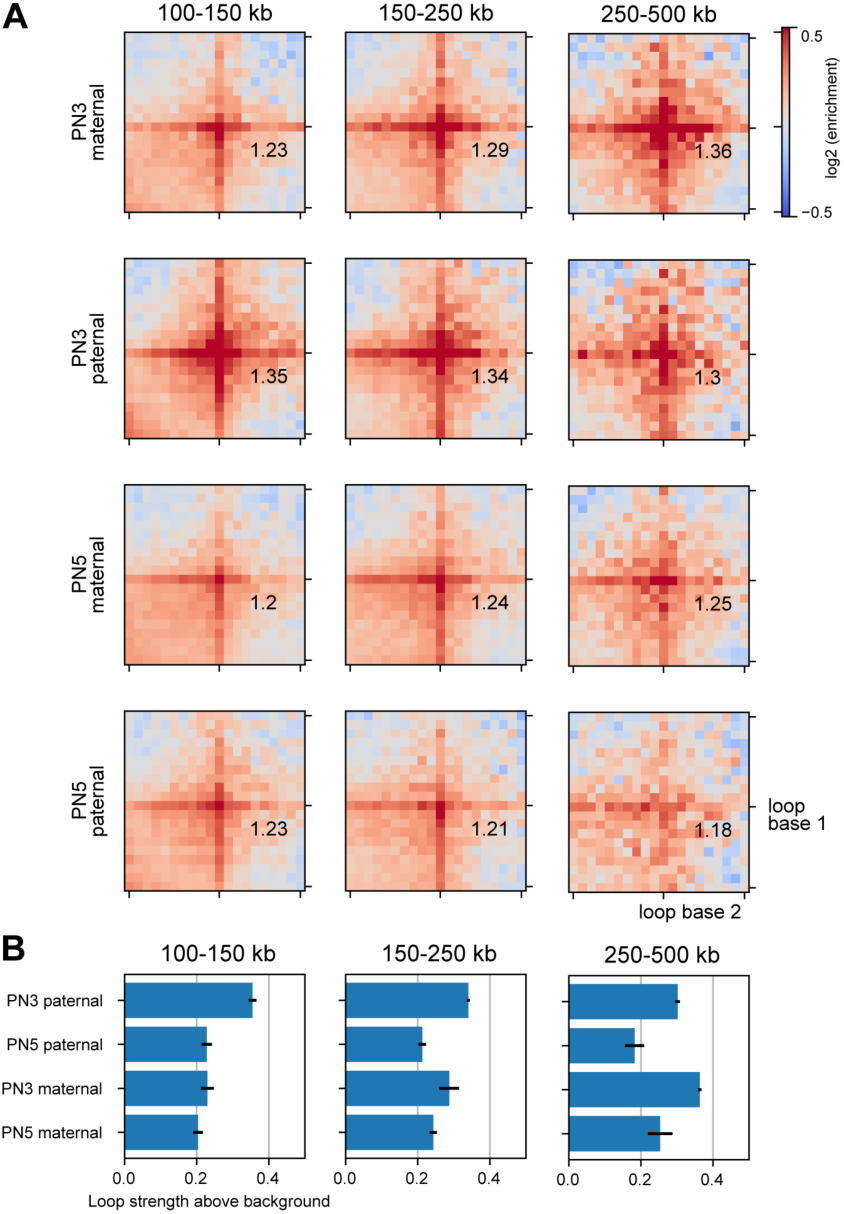
Zygotic paternal chromatin has stronger loops, Related to Figure 1. A) Average Hi-C loops, separated by distance, for the data re-analyzed from Du et al., 2017. Zygotic pronuclear stage 3 (PN3) and stage 5 correspond to S and G2 phases, respectively. The numeric values in each plot correspond to the fold enrichment in loop strength above background levels. Windows shown are a 190 kb region centered on the loop bases. B) Quantification of loop strength above background. Reported values are the fraction of enrichment above background. Error bars displayed are the 95% confidence intervals obtained by bootstrapping the experimental replicates.

**Supplementary Figure 2:**
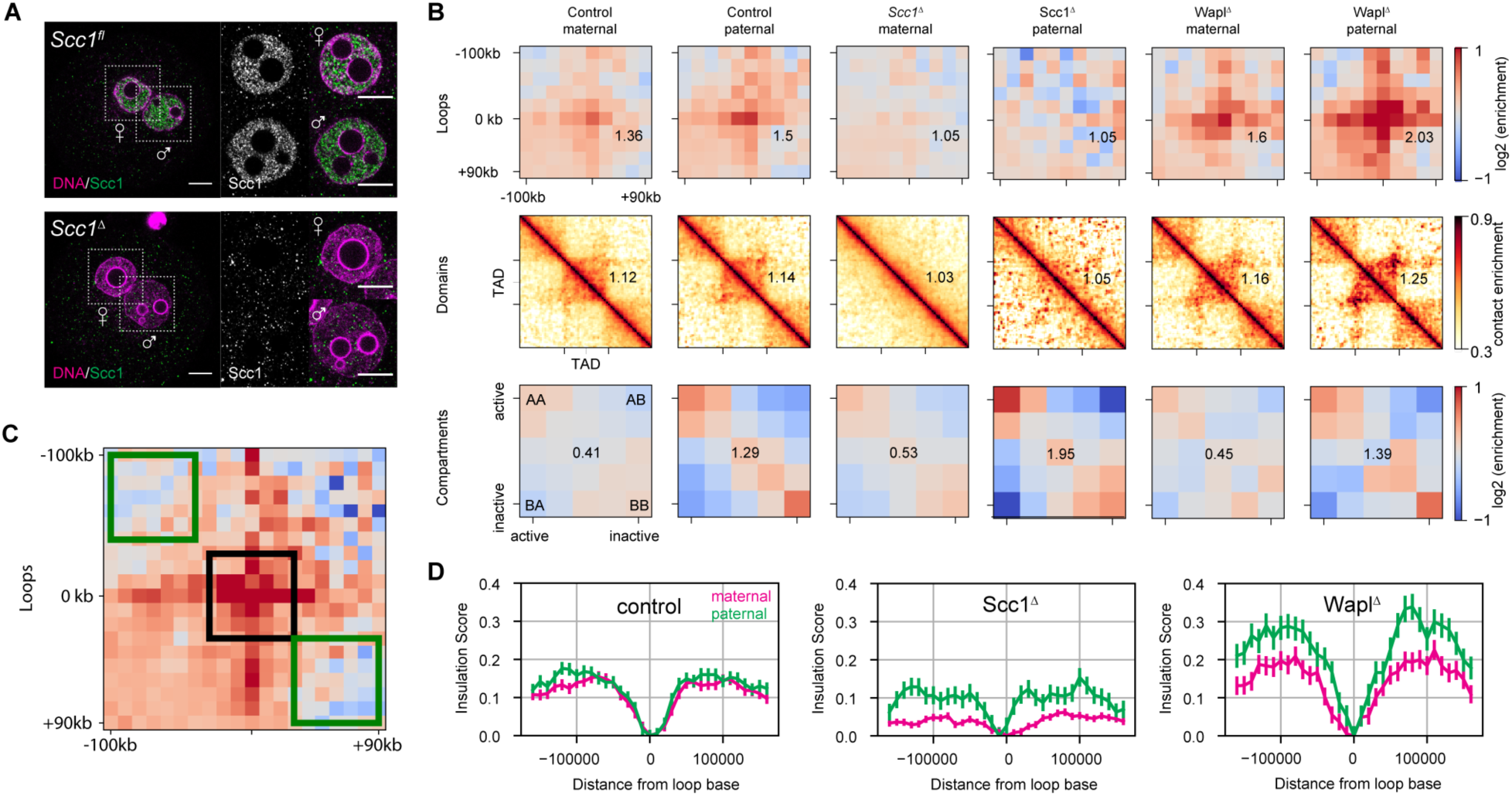
Additional information on conditional knockouts, Related to Figure 2. A) Immunofluorescence staining of Scc1 in *in situ* fixed *Scc1*^*fl*^ (n=15) and *Scc1*^*Δ*^ zygotes (n=12). DNA in magenta, Scc1 in grey/green. Images were adjusted in brightness/contrast in the individual channels using ImageJ. Scale bar: 10μm. Left: Single z-slice of zygotes. Right: Single z-slice of the maximum cross-section area of maternal and paternal nuclei. Cropped area is indicated. B) Loops, TADs, and compartment saddle plots for the control, *Scc1*^*Δ*^ and *Wapl*^*Δ*^ conditions shown separately for the maternal and paternal data. C) Loop strengths were calculated using the three 60 x 60 kb square regions shown. The average value within the middle box (black) was divided by the average of the combined top left and bottom right (green) boxes. The resulting number was subtracted by 1 to indicate the fractional increase in loop strength above the background. D) Insulation scores calculated with a diamond of size 40 kb, with the “zero” position denoting a domain boundary identified previously in CH12-LX cells (Rao et. al, 2014).

**Supplementary Figure 3:**
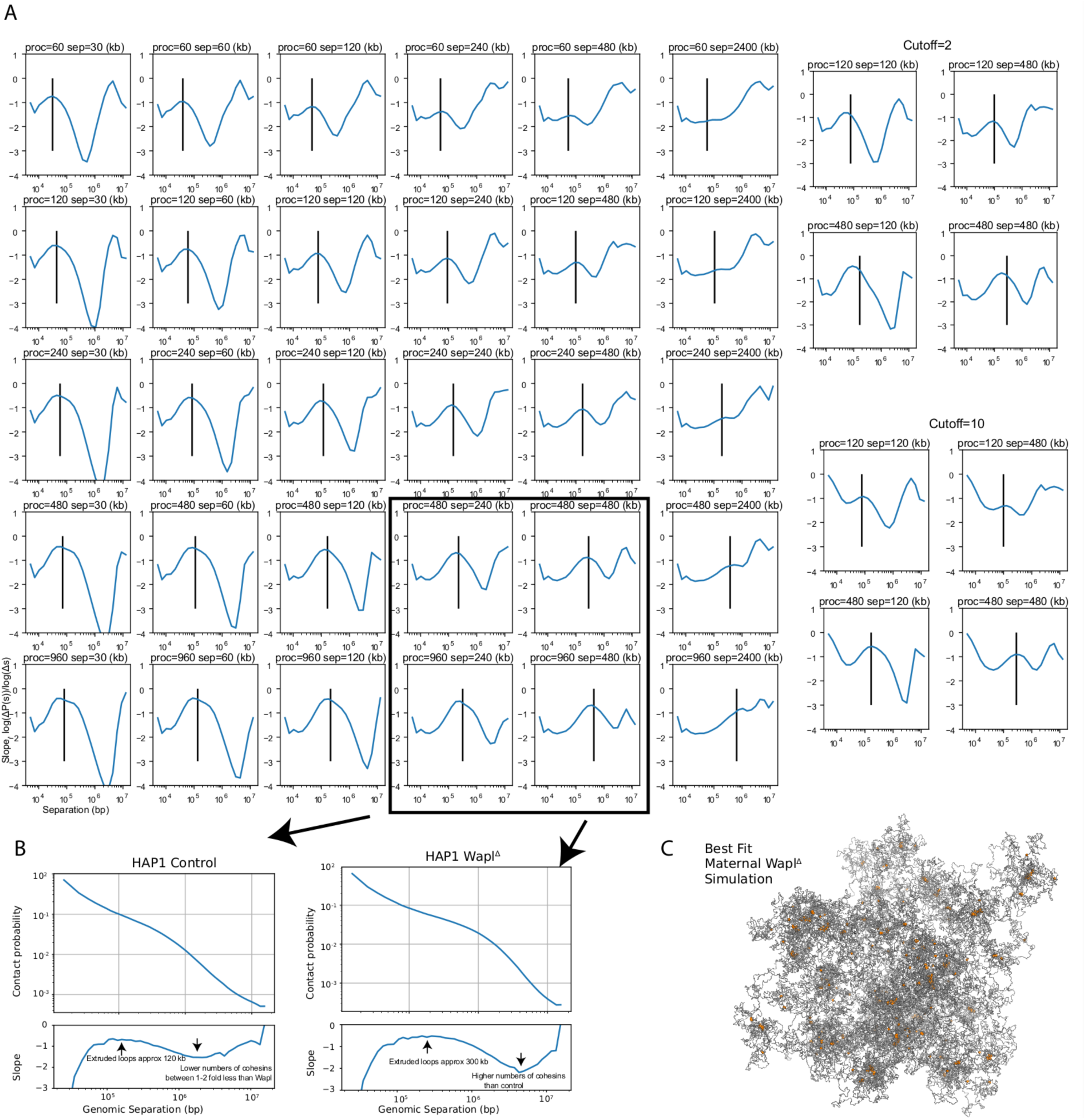
P_c_(s) curves, Related to Figure 3. A) Slopes of P_c_(s) curves as a function of genomic separation for N=30000 polymer models with loop extrusion. The rows show different loop extrusion processivities (proc), and columns show different linear separations (sep) between cohesins; the latter is related to the number of cohesins via the relation: separations = (chromosome length) / (number of bound cohesins). Vertical line on each plot indicates the average extruded loop length. All P_c_(s) plots in the left 6 rows were calculated for a Hi-C contact radius of 5 monomers (75 nm). Plots on the right are a subset of plots on the left, for contact radius of 2 monomers (30 nm), and 10 monomers (150 nm), indicating that the inferred average extruded loop length does not vary significantly with the choice of Hi-C capture radius. Note that average extruded loop length is different from processivity, especially in a dense regime where processivity is greater than separation; due to stalling of cohesins when encountering each other and at simulated TAD boundaries, the average loop length then becomes less than processivity; see Goloborodko et al., 2016 for details. B) The analysis of the slope of log(P_c_(s)) applied on recently published *Wapl*^*Δ*^ Hi-C data (see Haarhuis et al., 2017). Consistently with experimental FRAP data (Haarhuis et al., 2017), we find that in *Wapl*^*Δ*^ conditions the processivity, which is linearly related to the chromatin-bound lifetime of cohesin, is increased >2 above control conditions. Similarly, we find that the numbers of bound cohesins is >1 but less than 2-fold enriched above controls in *Wapl*^*Δ*^; this is consistent with quantitative western blot data, showing a 1.5-fold enrichment for cohesins in *Wapl*^*Δ*^ versus controls (Haarhuis et al., 2017). C) A representative image of the maternal *Wapl*^*Δ*^ simulation. The chromatin fiber is coloured in gray, and the locations of the cohesins coloured in orange, indicating that some cohesin vermicelli is visibly formed.

**Supplementary Figure 4:**
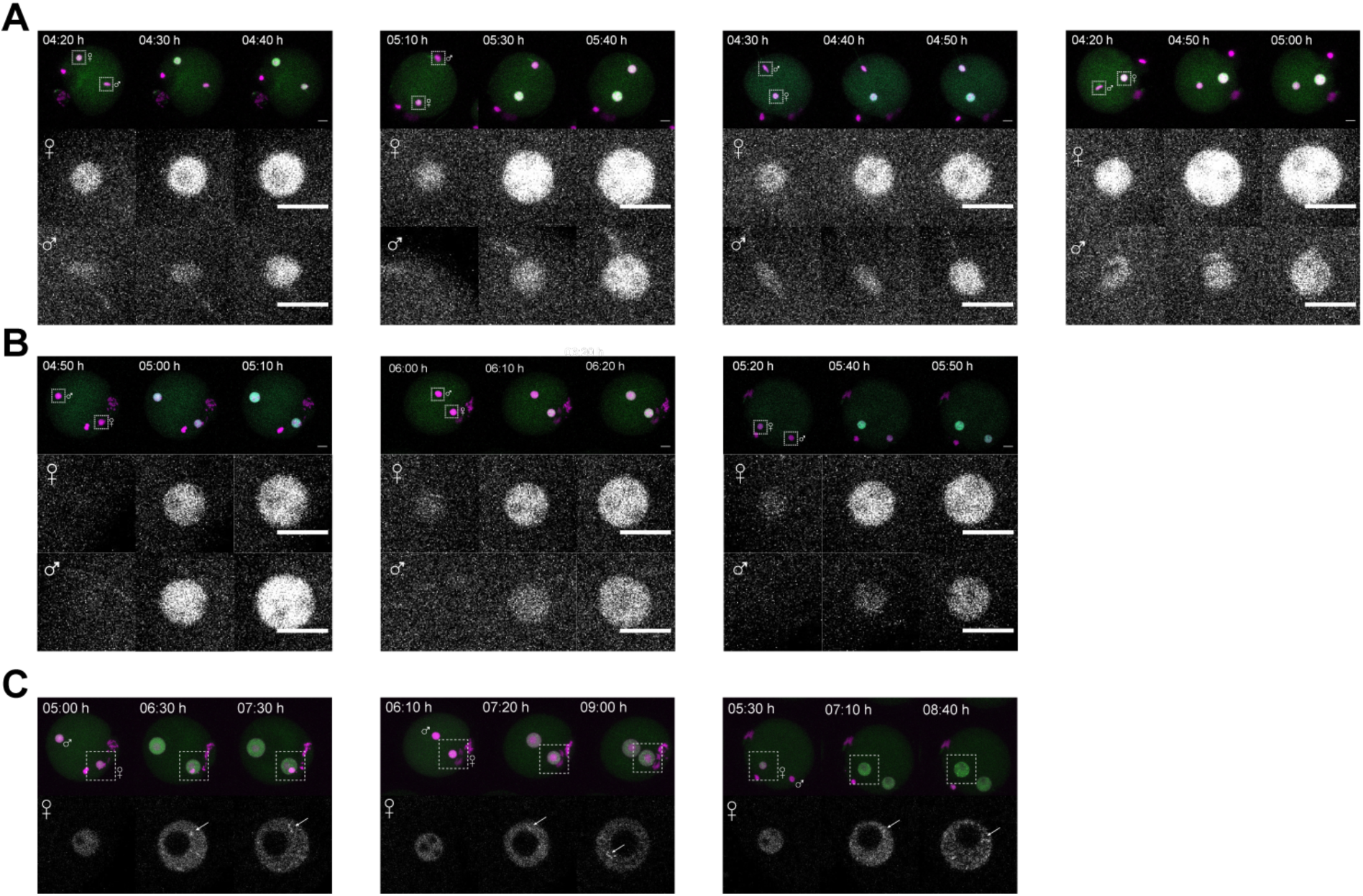
Live-cell imaging of wildtype and *Scc1ΔWaplΔ* zygotes expressing Scc1-EGFP and H2B-mCherry, Related to Figure 4. A) Onset of Scc1-EGFP accumulation in nuclei of wildtype zygotes (n=4). Top: Z-stack maximum intensity projection of whole zygotes. Bottom: Z-stack maximum intensity projection of maternal and paternal nuclei. B) Onset of Scc1-EGFP accumulation in nuclei of *Scc1*^*Δ*^*Wapl*^*Δ*^ zygotes (n=3). Top: Z-stack maximum intensity projection of whole zygotes. Bottom: Z-stack maximum intensity projection of maternal and paternal nuclei. C) Onset of vermicelli formation in *Scc1*^*Δ*^*Wapl*^*Δ*^ zygotes (n=3) corresponding to B. Top: Z-stack maximum intensity projection of whole zygotes. Bottom: Single z-slices of maternal nuclei.

**Supplementary Figure 5:**
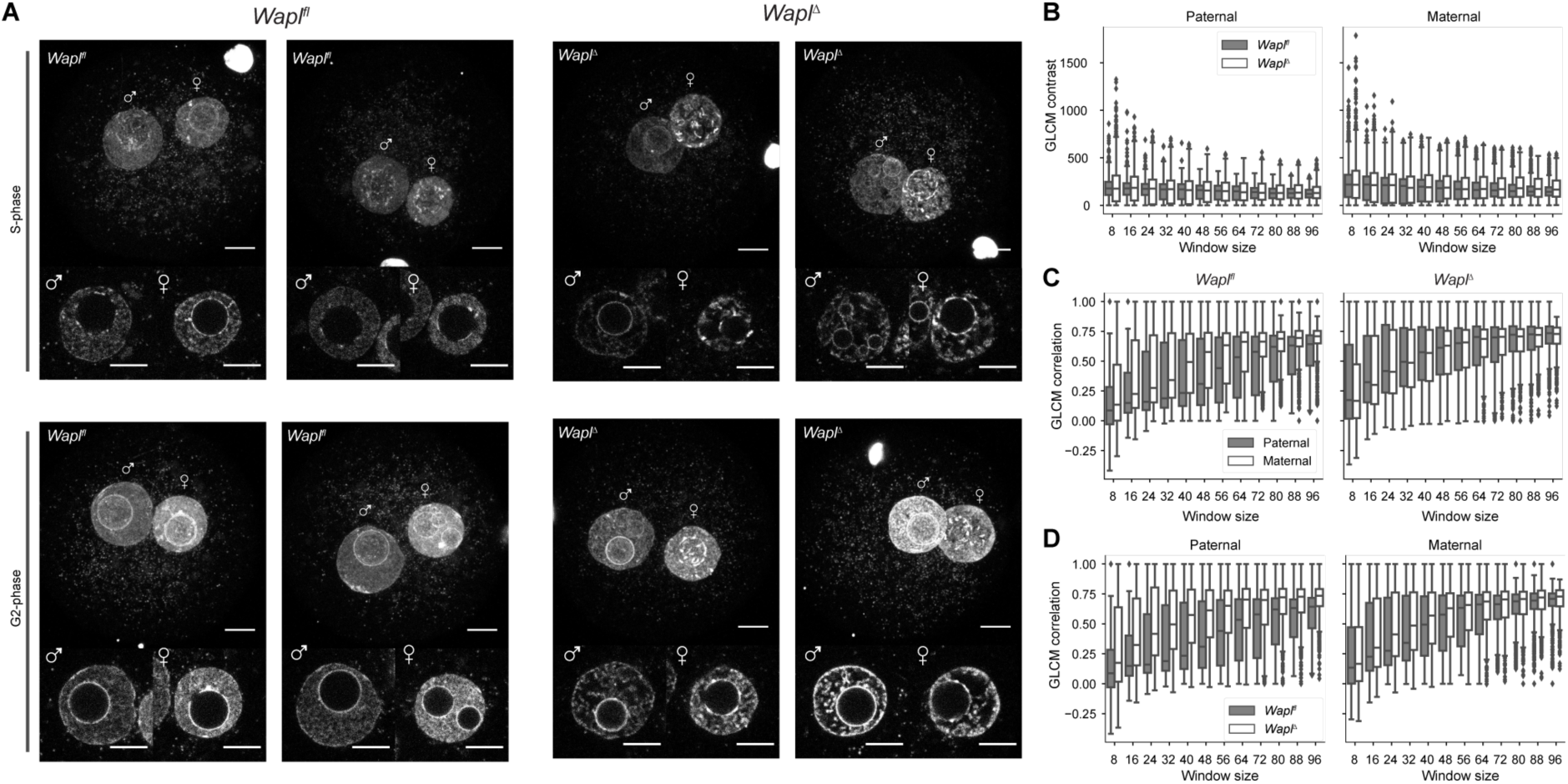
DNA staining of *Wapl*^*fl*^ and *Wapl*^*Δ*^ zygotes, Related to Figure 5. A) Representative images of fixed *Wapl*^*fl*^ and *Wapl*^*Δ*^ zygotes stained with DAPI. Zygotes were collected during S phase (9 h 45 min post fertilization; n=7 *Wapl*^*fl*^, n=15 *Wapl*^*Δ*^) or G2 phase (14 h post-fertilization; n=3 *Wapl*^*fl*^, n=8 *Wapl*^*Δ*^). Top: Z-stack maximum intensity projection (MIP) of zygotes. Bottom: Individual maternal and paternal nuclei. Settings were adjusted for z-slices and MIPs individually, but in the same manner for *Wapl*^*fl*^ and *Wapl*^*Δ*^ zygotes. Scale bar: 10μm. B) Boxplots showing GLCM contrast (local variation of intensity) in paternal and maternal nuclei of *Wapl*^*fl*^ (grey, n=10) and *Wapl*^*Δ*^ (white, n=12) zygotes with increasing window sizes. C) Boxplots showing GLCM correlation (linear dependence of intensity between adjacent pixels) in paternal (grey) and maternal (white) nuclei in *Wapl*^*fl*^ (n=10) and *Wapl*^*Δ*^ (n=12) zygotes with increasing window sizes. D) Boxplots showing GLCM correlation (linear dependence of intensity between adjacent pixels) in paternal and maternal nuclei of *Wapl*^*fl*^ (grey, n=10) and *Wapl*^*Δ*^ (white, n=12) zygotes with increasing window sizes.

### ACKNOWLEDGEMENTS

We thank M. Coelho Correia da Silva and M. H. Idarraga-Amado for providing conditional *Wapl* mice and reagents, and A. Hirsch, K. Klien and N. Laumann-Lipp for technical assistance with pronuclear extractions. We are grateful to J. Nuebler for discussions and suggestions for simulations and sharing codes, and to A. Goloborodko for suggesting the derivative-based analyses of P_c_(s) curves. Illumina sequencing was performed at the VBCF NGS Unit (<u>http://www.vbcf.ac.at</u>). We thank the staff of Vienna Biocenter BioOptics facility for assistance with imaging and analysis. J.G. is an associated student of the DK Chromosome Dynamics supported by the grant W1238-B20 from the Austrian Science Fund (FWF) and supported by the European Research Council.. H.B.B. is grateful for support by the Natural Sciences and Engineering Research Council of Canada, PGS-D. I.M.F. is grateful for support from the Darwin Trust of Edinburgh. W.A.B. is supported by the UK Medical Research Council. This work was funded by the Austrian Academy of Sciences and by the European Research Council (ERC-StG-336460 ChromHeritance) to K.T.-K.. The work in the Mirny laboratory is supported by R01 GM114190, U54 DK107980 from the National Institute of Health, and 1504942 from the National Science Foundation.

## AUTHOR CONTRIBUTIONS

J.G. and H.B.B. contributed equally. K.T.-K. conceived the project. J.G. supervised by K.T.-K. performed snHi-C and imaging experiments. H.B.B. supervised by L.A.M. developed and performed snHi-C data analysis and simulations. M.I. performed simulations. I.M.F. contributed expertise and performed image analysis. S.L. provided novel mouse strains and imaging reagents. J.M.P. provided a novel mouse strain. H.B.B., J.G., M.I. and I.M.F. prepared the figures. H.B.B., J.G., M.I., I.M.F., W.A.B., L.A.M. and K.T.-K. wrote the manuscript with input from all authors.

### Contact for reagent and resource sharing

Further information and requests for resources and reagents should be directed to and will be fulfilled by the Lead Contact, Kikue Tachibana-Konwalski.

## METHODS

### Mice

The care and use of the mice were carried out in agreement with the authorizing committee according to the Austrian Animal Welfare law and the guidelines of the International guiding principles for biomedical research involving animals (CIOMS, the Council for International Organizations of Medical Sciences). Mice were kept at a daily cycle of 14 hours light and 10 hours dark with access to food *ad libitum*. All mice were bred in the IMBA animal facility. *Scc1*^*fl/fl*^ mice were bred on a mixed background (B6, 129, Sv). Wapl ^*fl/fl*^ mice were bred on a primarily C57BL/6J background. *Scc1*^*fl/fl*^ *Wapl*^*fl/fl*^ mice were bred on the same mixed background as *Scc1* ^*fl/fl*^ mice. Experimental mice were obtained by mating of homozygous floxed females to homozygous floxed males carrying *Tg(Zp3-Cre)* (Lewandoski *et al*, 1997; Lan *et al*, 2004). To obtain zygotes B6CBAF1 stud males were mated to *Scc1* ^*fl/fl*^ *Tg(Zp3-Cre)*, while C57BL/6J stud males were used for mating *Wapl* ^*fl/fl*^ *Tg(Zp3-Cre)* females. Sperm for *in vitro* fertilization of *Scc1* ^*fl/fl*^ *Wapl* ^*fl/fl*^ *Tg(Zp3-Cre*) oocytes was obtained from B6CBAF1 stud males.

### Zygote collection

To obtain zygotes 3-5 week old female mice were superovulated by intraperitoneal injection of PMSG (pregnant mare’s serum gonadotropin; 5 IU, Folligon, Intervet or 5 IU, Prospecbio) followed by hCG (human chorionic gonadotropin; 5 IU, Chorulon, Intervet) injection 48 hours later. Females were mated to wildtype stud males overnight. The following morning zygotes were released from the ampullae and treated with hyaluronidase to remove cumulus cells.

### Single-nucleus Hi-C

Single-nucleus Hi-C was carried out as described before (Flyamer et al., 2017). After pronuclear extraction *Scc1*^*fl/fl*^ *Tg(Zp3-Cre)* pronuclei used in the experiments were fixed around 19-22 hours post hCG injection (corresponding to about 7-10 hours post fertilization) and therefore are expected to be in G1/S-phase of the cell cycle. *Wapl*^*fl/fl*^ *Tg(Zp3-Cre)* were fixed later around 23-27.5 hours post hCG injection (corresponding to about 11-15.5 hours post fertilization) and are expected to be in S/G2 phase of the cell cycle. To obtain G2 phase data zygotes were fixed 27 hours post hCG injection (corresponding to about 15 hours post fertilization) and lysed, pronuclei were separated into different wells after SDS lysis according to their size. No blinding or randomization were used for handling of the cells.

Briefly, after pronuclei were isolated, they were fixed in 2% formaldehyde for 15 minutes, then lysed on ice in lysis buffer (10 mM Tris-HCl pH 8.0, 10 mM NaCl, 0.5% (v/v) NP-40 substitute (Sigma), 1% (v/v) Triton X-100 (Sigma), 1× Halt^TM^ Protease Inhibitor Cocktail (Thermo Scientific)) for at least 15 minutes. Then the pronuclei were washed once through PBS and 1x NEB3 buffer (NEB) with 0.6% SDS, in which they were then incubated at 37° for 2 hours with shaking in humidified atmosphere. Then pronuclei were washed once in 1x DpnII buffer (NEB) with 2x BSA (NEB), and then chromatin was digested overnight in 9ul of the same solution but with 5 U DpnII (NEB). The nuclei were then washed once through PBS, then through 1xT4 ligase buffer (50 mM Tris-HCl, 10 mM MgCl_2_, 1 mM ATP, 10 mM DTT, pH 7.5). Then the nuclei were incubated in the same buffer but with 5U T4 DNA ligase (Thermo Scientific) at 16° with 50 rpm rotation for 4.5 hours, and then for 30 min at room temperature. Whole-genome amplification was performed using illustra GenomiPhi v2 DNA amplification kit (GE Healthcare) with decrosslinking nuclei at 65° overnight in sample buffer. High molecular weight DNA was purified using AMPure XP beads (Beckman Coulter), and 1 ug was used to prepare Illumina libraries for sequencing (by VBCF NGS Unit, csf.ac.at) after sonicating to ∼300-1300 bp. Libraries were sequenced on HiSeq 2500 v4 with 125 bp paired end reads, between 10 and 24 cells per lane.

### DNA and Scc1 staining

After removal of cumulus cells by hyaluronidase treatment zygotes were fixed in 4% PFA for 30 min, before permeabilization in 0.2% Triton X-100/PBS (PBSTX) for 30 min. Cells were then blocked in 10% goat serum (Dako) in PBSTX either at 4C overnight or for several hours at 4C followed by room temperature incubation. Cells were incubated overnight at 4C in primary antibody (anti-Scc1, Millipore #05-908, 1:250). After washing in blocking solution for at least 30 min, incubation with the secondary antibody (anti-mouse IgG (H+L), Thermo Fisher Scientific #A-11001, 1:500) was carried out for 1h at room temperature. Another set of washing steps in 0.2% Triton X-100/PBS was followed by a quick PBS wash and mounting of the cells in Vectashield containing DAPI (Vector labs) using imaging spacers (Sigma Aldrich). *In situ* fixed zygotes were imaged on a confocal microscope (LSM780, Zeiss) using a 63x, 1.4NA oil objective. Presence of DNA compaction reminiscent of vermicelli in Wapl zygotes was classified using ImageJ and 3D visualization by Imaris (8.1.2). No blinding or randomization were used for handling of the cells.

### Live cell imaging

*In vitro* fertilization after *in vitro* maturation was performed as described before (Ladstätter and Tachibana-Konwalski, 2017). Oocytes from 2-5 month old females were isolated by puncturing of ovaries with hypodermic needles in the presence of 0.2 mM IBMX, 20% FBS (Gibco) and 6 mg/ml fetuin (Sigma Aldrich). After microinjection of oocytes with H2B-mCherry (0.5 pmol; 187 ng/μl) and Scc1-EGFP (0.4 pmol; 260 ng/μl), oocytes were cultured for 1-1.5 h and then released from IBMX inhibition by washing in M16. Following *in vitro* maturation in the incubator (low oxygen conditions: 5% CO2, 5% O2, 90% N2; 37°C), cells were scored for extrusion of the first polar body and MII eggs were *in vitro* fertilized 10-11 hours post release from IBMX inhibition. The sperm was obtained from the *cauda epididimis* and *vas deferens* of B6CBAF1 stud males and was capacitated in fertilization medium (Cook) in a tilted cell culture dish for at least 30 min. Motile sperm from the surface of the dish was used for *in vitro* fertilization of the *in vitro* maturated eggs. After 3 h zygotes were washed in M16 and imaged. Live-cell imaging of zygotes microinjected with fluorescent fusion proteins was performed on a confocal microscope (LSM 800, Zeiss) equipped with an incubation chamber suited for live-cell imaging (5% CO2, 37°C). Zygotes were kept in ∼3μl cleavage medium (Research Vitro Cleave, Cooks Austria GmbH) under mineral oil (Sigma Aldrich) for the duration of the imaging. Movies were taken using a 63x, 1.20NA water immersion objective, taking 25 z-slices (48μm) every 10 minutes.

### snHi-C data analysis

snHi-C data were processed similarly as in (Flyamer et al., 2017). Briefly, reads were mapped to the mm9 genome using *hiclib* (which applies iterative mapping with *bowtie2*) and then filtered. These data were then converted into *Cooler* files with heatmaps at different resolutions for downstream analysis.

We applied the same methods for quantification of different features of spatial organization of the genome as previously (Flyamer et al., 2017). Briefly, we used GC-content as a proxy for A/B compartmentalization signal and constructed 5x5 percentile-binned matrices to quantify strength of compartment segregation. For average analysis of TADs we used published TAD coordinates (Rao et al., 2014) for the CH12-LX mouse cell line. We averaged Hi-C maps of all TADs and their neighbouring regions, chosen to be of the same length as the TAD, after rescaling each TAD to a 90x90 matrix. For visualization, the contact probability of these matrices was rescaled to follow a shallow power law with distance (-0.25 scaling). Similarly, we analyzed loops by summing up snHi-C contact frequencies for loop coordinates identified in Rao et al., 2014 for CH12-LX mouse cells. By averaging 20x20 matrices surrounding the loops and dividing the final result by similarly averaged control matrices, we removed the effects of distance-dependence. Control loop matrices were obtained by averaging 20x20 matrices centered on the locations of randomly shifted positions of known loops; shifts ranged from 100 to 1100 Kb with 100 shifts for each loop. For display and visual consistency with the loop strength quantification, we set the background levels of interaction to 1; the background is defined as the green boxes in **Figure S2C** described below.

For the quantification of loop strength, we divided the average signal in the middle 6x6 submatrix by the average signal in top-left and bottom-right (at the same distance from the main diagonal) 6x6 submatrices (see **Figure S2C**). To obtain the 95% confidence intervals on the loop strengths we applied bootstrapping: using the pooled single cell data, we randomly sampled N loops with replacement (where N equals the total number of loops used in the original samples), and calculated the loop strengths from this random sample. We performed this procedure 10,000 times for each condition, using the sorted set of 10,000 strength values to obtain the confidence intervals.

Contact probability, P_c_(s), curves were computed from 10 kb binned snHi-C data. We divided the linear genomic separations into logarithmic bins with a factor of 1.3. Data within these log-spaced bins (at distance, s) were averaged to produce the value of P_c_(s).

### DAPI texture analysis

All analysis was performed using python (3.5.2) scientific stack (numpy-1.13.1, scipy-0.19.1, pandas-0.20.2, matplotlib-2.0.2) with image analysis specific functions from scikit-image-0.13.0. Images of zygotes stained with DAPI were automatically thresholded plane-by-plane using the Otsu method after median filtering (using 5x5 pixel square) and gaussian blurring with sigma 5. After binary closing and removal of small holes (<50 pixels), elongated (major axis length >1.5 times longer than minor axis) and misshapen (circularity below 0.5) holes were also removed. Then objects with area below 5000 pixels or above 100,000 pixels were removed, along with dim (with average intensity below double average intensity of the whole image) and misshapen (circularity below 0.15) objects. Then all planes were combined to form a single object annotation for the whole z-stack, and again small (<500 pixels in volume) and large (>100,000 pixels in volume) objects were removed.

For the coefficient of variation analysis, all pixels from each object the segmented image were used to calculate mean and standard deviation, then the CV from all objects in the image was averaged. For analysis with comparisons of maternal and paternal pronuclei, the images were additionally manually filtered to exclude improperly segmented ones. Then maternal and paternal origin of pronuclei was determined using their size: we considered paternal the ones where both the total volume and the biggest cross-section area were higher than in the other pronucleus, which we then considered maternal; we didn’t use the images where these two measurements disagreed.

For the GLCM analysis, we randomly generated 100 2D windows for each image, so that they are fully inside the segmented area, and moreover, so that if their size was increased by 14 pixels along x and y axis and they were shifted up or down by one z plane, they were still fully inside the masked region. We performed this for windows of sizes between 8×8 and 96×96 pixels, with each step increasing the sides of the windows by 8 pixels, in total 1200 windows per nucleus. We then applied GLCM analysis of correlation and contrast to each of the windows, and combined data from all windows together, recording their origin. P-values were calculated using the Mann-Whitney U test.

### Analysis of Du et al., 2017 data

Pre-processed, mapped valid pair files were obtained from GEO accession number GSE82185. These files were directly converted to the *Cooler* format (<u>https://github.com/mirnylab/cooler</u>) without any further filtering or processing using csort and cload functions. Averaging analysis for loops, TADs, compartments were performed as described previously (Flyamer et al., 2017) and summarized in the above section.

### Polymer simulations

Polymer simulations of loop extrusion were performed as in (Flyamer et al., 2017), but using updates to the simulation engine (Fudenberg et al., 2017). The simulation engine is build using the openmm-polymer package which relies on OpenMM-7 (Eastman et al., 2017). Parameters for simulations were as follows: 2000 MD steps per loop extrusion step. Simulations were performed either using N=30,000 monomers, or N=100,000 monomers. Simulations were initialized using a fractal globule or a mitotic chromosome model, as described in (Flyamer et al., 2017). Bi-directional TAD boundaries were placed at monomers 0, 1200, 1500, 2000,2900, 3900, 4300, 4800, 5600, 6100, 6500, 7600, 8300, 8900, 9500; and at positions shifted by multiples of 10,000 (10000, 11200, 11500, 12000, … 20000, 21200, 21500, 22000…). TAD boundaries were implemented as monomers that pause the loop extruding factor (LEF) translocation with probability 99.5 %. That would delay translocation of a LEF by on average 200 loop extrusion steps. All simulations were performed in periodic boundary conditions at a given density. For each simulation, we simulated 4000 steps of loop extrusion dynamics, starting with a random placement of LEFs at the beginning of a simulation.

We performed two types of simulations. A parameter sweep for processivity-separation values was performed for a system of 30,000 monomers for all pairwise combinations of the values of processivity of 60, 120, 240, 480, and 960kb, and the values of separation of 30, 60, 120, 240, 480, and 2400kb. The largest value of separation was to simulate 20-fold depletion of LEFs relative to WT model value of 120kb (Fudenberg et al., 2016). All simulations here were initialized with a 30,000 fragment of a mitotic chromosome model. We used density of 0.02 for these simulations.

A more complete simulation was performed using a system of 100,000 monomers, initialized from a mitotic chromosome model, or from a fractal globule for maternal and paternal chromosomes, respectively. Particular values of parameters were chosen based on a parameter sweep. We chose values of processivity and separation of 120kb for the control conditions model, the same values as used in (Fudenberg et al., 2016). For the model of SCC1 knock-out, we reduced the number of cohesins 20-fold, which corresponds to increasing separation to 2400kb. For the model of WAPL knock-out of maternal chromatin, we increased processivity 4-fold, but kept the separation at 120 kb. For WAPL knock-out of paternal chromatin, we best matched the difference in P_c_(s) in the s=100-500kb region by decreasing the processivity two-fold, but increasing separation by two-fold as compared to maternal. Additionally, to reflect the larger paternal pronuclear volume, we decreased the density of simulations two-fold, to 0.01.

We calculated P_c_(s) and simulated contact maps using contact radius of 5 monomers.

### Polymer simulation surface area and volume measurements

We calculated the surface area and volume of single polymer conformations using the MATLAB R2017a alphaShapes class. In brief, 3 polymer conformations were randomly sampled for each simulated chromatin condition from the system of 30,000 monomer simulated chromatin fibers. For a given configuration, we calculated the numbers of contacts in cis and trans using a cutoff radius of 5 monomer radii. Trans contacts were computed from the contacts of the polymer with its 26 periodic boundary images in theneighbouring simulation volumes. To calculate the surface area and volume, we defined spheres of radius 1 around each monomer using the “sphere” function, with input argument 10, and computed the alpha shape on the resulting set of points with alpha parameter 1.6 to account for variable bond distances between monomers due to the harmonic potential. The surface area and volumes were computed using the .surfaceArea and .volume methods respectively. Results of the polymer simulations were plotted against the calculated number of cis and trans contacts.

### Data and software availability

The snHi-C data has been deposited on NCBI GEO under the accession number: GSE100569. Polymer simulation code is available in the “examples” directory of the openmm-polymer library (<u>https://bitbucket.org/mirnylab/openmm-polymer</u>); analysis code of polymer configurations, including the surface area and volume measurements will be made available at the time of publication. snHi-C data processing code has been released as an example for the hiclib package (<u>https://bitbucket.org/mirnylab/hiclib</u>).

## REFERENCES

Adenot, P.G., Mercier, Y., Renard, J.P., and Thompson, E.M. (1997). Differential H4 acetylation of paternal and maternal chromatin precedes DNA replication and differential transcriptional activity in pronuclei of 1-cell mouse embryos. Development 124, 4615–4625.

Aoki, F., Worrad, D.M., and Schultz, R.M. (1997). Regulation of transcriptional activity during the first and second cell cycles in the preimplantation mouse embryo. Dev. Biol. 181, 296–307.

Bouniol, C. (1995). Endogenous Transcription Occurs at the 1-Cell Stage in the Mouse Embryo. Exp. Cell Res. 218, 57–62.

Burkhardt, S., Borsos, M., Szydlowska, A., Godwin, J., Williams, S.A., Cohen, P.E., Hirota, T., Saitou, M., and Tachibana-Konwalski, K. (2016). Chromosome Cohesion Established by Rec8-Cohesin in Fetal Oocytes Is Maintained without Detectable Turnover in Oocytes Arrested for Months in Mice. Curr. Biol. 26, 678–685.

Busslinger, G.A., Stocsits, R.R., van der Lelij, P., Axelsson, E., Tedeschi, A., Galjart, N., and Peters, J.-M. (2017). Cohesin is positioned in mammalian genomes by transcription, CTCF and Wapl. Nature 544, 503–507.

Ciosk, R., Shirayama, M., Shevchenko, A., Tanaka, T., Toth, A., Shevchenko, A., and Nasmyth, K. (2000). Cohesin’s Binding to Chromosomes Depends on a Separate Complex Consisting of Scc2 and Scc4 Proteins. Mol. Cell 5, 243–254.

Deardorff, M.A., Kaur, M., Yaeger, D., Rampuria, A., Korolev, S., Pie, J., Gil-Rodríguez, C., Arnedo, M., Loeys, B., Kline, A.D., et al. (2007). Mutations in Cohesin Complex Members SMC3 and SMC1A Cause a Mild Variant of Cornelia de Lange Syndrome with Predominant Mental Retardation. Am. J. Hum. Genet. 80, 485–494.

Dixon, J.R., Selvaraj, S., Yue, F., Kim, A., Li, Y., Shen, Y., Hu, M., Liu, J.S., and Ren, B. (2012). Topological domains in mammalian genomes identified by analysis of chromatin interactions. Nature 485, 376–380.

Du, Z., Zheng, H., Huang, B., Ma, R., Wu, J., Zhang, X., He, J., Xiang, Y., Wang, Q., Li, Y., et al. (2017). Allelic reprogramming of 3D chromatin architecture during early mammalian development. Nature 547, 232–235.

Eastman, P., Swails, J., Chodera, J.D., McGibbon, R.T., Zhao, Y., Beauchamp, K.A., Wang, L.- P., Simmonett, A.C., Harrigan, M.P., Stern, C.D., et al. (2017). OpenMM 7: Rapid development of high performance algorithms for molecular dynamics. PLOS Comput. Biol. 13, e1005659.

Flavahan, W.A., Drier, Y., Liau, B.B., Gillespie, S.M., Venteicher, A.S., Stemmer-Rachamimov, A.O., Suvà, M.L., and Bernstein, B.E. (2015). Insulator dysfunction and oncogene activation in IDH mutant gliomas. Nature 529, 110–114.

Flyamer, I.M., Gassler, J., Imakaev, M., Brandão, H.B., Ulianov, S. V., Abdennur, N., Razin, S. V., Mirny, L.A., and Tachibana-Konwalski, K. (2017). Single-nucleus Hi-C reveals unique chromatin reorganization at oocyte-to-zygote transition. Nature 544, 110–114.

Franke, M., Ibrahim, D.M., Andrey, G., Schwarzer, W., Heinrich, V., Schöpflin, R., Kraft, K., Kempfer, R., Jerković, I., Chan, W.-L., et al. (2016). Formation of new chromatin domains determines pathogenicity of genomic duplications. Nature 538, 265–269.

Fudenberg, G., and Imakaev, M. (2017). FISH-ing for captured contacts: towards reconciling FISH and 3C. Nat. Methods 14, 673–678.

Fudenberg, G., Imakaev, M., Lu, C., Goloborodko, A., Abdennur, N., and Mirny, L.A. (2016). Formation of Chromosomal Domains by Loop Extrusion. Cell Rep. 15, 2038–2049.

Gandhi, R., Gillespie, P.J., and Hirano, T. (2006). Human Wapl Is a Cohesin-Binding Protein that Promotes Sister-Chromatid Resolution in Mitotic Prophase. Curr. Biol. 16, 2406–2417.

Giorgetti, L., Lajoie, B.R., Carter, A.C., Attia, M., Zhan, Y., Xu, J., Chen, C.J., Kaplan, N., Chang, H.Y., Heard, E., et al. (2016). Structural organization of the inactive X chromosome in the mouse. Nature 535, 575–579.

Haarhuis, J.H.I., van der Weide, R.H., Blomen, V.A., Yáñez-Cuna, J.O., Amendola, M., van Ruiten, M.S., Krijger, P.H.L., Teunissen, H., Medema, R.H., van Steensel, B., et al. (2017). The Cohesin Release Factor WAPL Restricts Chromatin Loop Extension. Cell 169, 693–707.e14.

Halverson, J.D., Smrek, J., Kremer, K., and Grosberg, A.Y. (2014). From a melt of rings to chromosome territories: the role of topological constraints in genome folding. Rep. Prog. Phys. 77, 22601.

Hamatani, T., Carter, M.G., Sharov, A.A., and Ko, M.S.H. (2004). Dynamics of Global Gene Expression Changes during Mouse Preimplantation Development. Dev. Cell 6, 117–131.

van der Heijden, G.W., Dieker, J.W., Derijck, A.A.H.A., Muller, S., Berden, J.H.M., Braat, D.D.M., van der Vlag, J., and de Boer, P. (2005). Asymmetry in Histone H3 variants and lysine methylation between paternal and maternal chromatin of the early mouse zygote. Mech. Dev. 122, 1008–1022.

Hug, C.B., Grimaldi, A.G., Kruse, K., and Vaquerizas, J.M. (2017). Chromatin Architecture Emerges during Zygotic Genome Activation Independent of Transcription. Cell 169, 216– 228.e19.

Inoue, A., Jiang, L., Lu, F., Suzuki, T., and Zhang, Y. (2017). Maternal H3K27me3 controls DNA methylation-independent imprinting. Nature 547, 419–424.

Kagey, M.H., Newman, J.J., Bilodeau, S., Zhan, Y., Orlando, D.A., van Berkum, N.L., Ebmeier, C.C., Goossens, J., Rahl, P.B., Levine, S.S., et al. (2010). Mediator and cohesin connect gene expression and chromatin architecture. Nature 467, 430–435.

Ke, Y., Xu, Y., Chen, X., Feng, S., Liu, Z., Sun, Y., Yao, X., Li, F., Zhu, W., Gao, L., et al. (2017). 3D Chromatin Structures of Mature Gametes and Structural Reprogramming during Mammalian Embryogenesis. Cell 170, 367–381.e20.

Krantz, I.D., McCallum, J., DeScipio, C., Kaur, M., Gillis, L.A., Yaeger, D., Jukofsky, L., Wasserman, N., Bottani, A., Morris, C.A., et al. (2004). Cornelia de Lange syndrome is caused by mutations in NIPBL, the human homolog of Drosophila melanogaster Nipped-B. Nat. Genet. 36, 631–635.

Kueng, S., Hegemann, B., Peters, B.H., Lipp, J.J., Schleiffer, A., Mechtler, K., and Peters, J.-M. (2006). Wapl Controls the Dynamic Association of Cohesin with Chromatin. Cell 127, 955–967.

Ladstätter, S., and Tachibana-Konwalski, K. (2016). A Surveillance Mechanism Ensures Repair of DNA Lesions during Zygotic Reprogramming. Cell 167, 1774–1787.e13.

Lan, Z.-J., Xu, X., and Cooney, A.J. (2004). Differential Oocyte-Specific Expression of Cre Recombinase Activity in GDF-9-iCre, Zp3cre, and Msx2Cre Transgenic Mice1. Biol. Reprod. 71, 1469–1474.

Larson, A.G., Elnatan, D., Keenen, M.M., Trnka, M.J., Johnston, J.B., Burlingame, A.L., Agard, D.A., Redding, S., and Narlikar, G.J. (2017). Liquid droplet formation by HP1a suggests a role for phase separation in heterochromatin. Nature 547, 236–240.

Lewandoski, M., Wassarman, K.M., and Martin, G.R. (1997). Zp3–cre, a transgenic mouse line for the activation or inactivation of loxP-flanked target genes specifically in the female germ line. Curr. Biol. 7, 148–151.

Lopez-Serra, L., Lengronne, A., Borges, V., Kelly, G., and Uhlmann, F. (2013). Budding yeast Wapl controls sister chromatid cohesion maintenance and chromosome condensation. Curr. Biol. 23, 64–69.

Lupiáñez, D.G., Kraft, K., Heinrich, V., Krawitz, P., Brancati, F., Klopocki, E., Horn, D., Kayserili, H., Opitz, J.M., Laxova, R., et al. (2015). Disruptions of Topological Chromatin Domains Cause Pathogenic Rewiring of Gene-Enhancer Interactions. Cell 161, 1012–1025.

Mayer, W., Niveleau, A., Walter, J., Fundele, R., and Haaf, T. (2000). Demethylation of the zygotic paternal genome. Nature 403, 501–502.

Mizuguchi, T., Fudenberg, G., Mehta, S., Belton, J.-M., Taneja, N., Folco, H.D., FitzGerald, P., Dekker, J., Mirny, L., Barrowman, J., et al. (2014). Cohesin-dependent globules and heterochromatin shape 3D genome architecture in S. pombe. Nature 516, 432–435.

Musio, A., Selicorni, A., Focarelli, M.L., Gervasini, C., Milani, D., Russo, S., Vezzoni, P., and Larizza, L. (2006). X-linked Cornelia de Lange syndrome owing to SMC1L1 mutations. Nat. Genet. 38, 528–530.

Nagano, T., Lubling, Y., Várnai, C., Dudley, C., Leung, W., Baran, Y., Mendelson Cohen, N., Wingett, S., Fraser, P., and Tanay, A. (2017). Cell-cycle dynamics of chromosomal organization at single-cell resolution. Nature 547, 61–67.

Naumova, N., Imakaev, M., Fudenberg, G., Zhan, Y., Lajoie, B.R., Mirny, L. a, and Dekker, J. (2013). Organization of the mitotic chromosome. Science 342, 948–953.

Nolen, L.D., Boyle, S., Ansari, M., Pritchard, E., and Bickmore, W.A. (2013). Regional chromatin decompaction in Cornelia de Lange syndrome associated with NIPBL disruption can be uncoupled from cohesin and CTCF. Hum. Mol. Genet. 22, 4180–4193.

Nora, E.P., Lajoie, B.R., Schulz, E.G., Giorgetti, L., Okamoto, I., Servant, N., Piolot, T., van Berkum, N.L., Meisig, J., Sedat, J., et al. (2012). Spatial partitioning of the regulatory landscape of the X-inactivation centre. Nature 485, 381–385.

Nora, E.P., Goloborodko, A., Valton, A.-L., Gibcus, J.H., Uebersohn, A., Abdennur, N., Dekker, J., Mirny, L.A., and Bruneau, B.G. (2017). Targeted Degradation of CTCF Decouples Local Insulation of Chromosome Domains from Genomic Compartmentalization. Cell 169, 930– 944.e22.

Oswald, J., Engemann, S., Lane, N., and Mayer, W. (2000). Active demethylation of the paternal genome in the mouse zygote. Curr. Biol. 475–478.

Parelho, V., Hadjur, S., Spivakov, M., Leleu, M., Sauer, S., Gregson, H.C., Jarmuz, A., Canzonetta, C., Webster, Z., Nesterova, T., et al. (2008). Cohesins Functionally Associate with CTCF on Mammalian Chromosome Arms. Cell 132, 422–433.

Peters, J.-M., Tedeschi, A., and Schmitz, J. (2008). The cohesin complex and its roles in chromosome biology. Genes Dev. 22, 3089–3114.

Phillips-Cremins, J.E., Sauria, M.E.G., Sanyal, A., Gerasimova, T.I., Lajoie, B.R., Bell, J.S.K., Ong, C.-T., Hookway, T.A., Guo, C., Sun, Y., et al. (2013). Architectural Protein Subclasses Shape 3D Organization of Genomes during Lineage Commitment. Cell 153, 1281–1295.

Rao, S., Huang, S.-C., Glenn St. Hilaire, B., Engreitz, J.M., Perez, E.M., Kieffer-Kwon, K.-R., Sanborn, A.L., Johnstone, S.E., Bochkov, I.D., Huang, X., et al. (2017). Cohesin Loss Eliminates All Loop Domains, Leading To Links Among Superenhancers And Downregulation Of Nearby Genes. bioRxiv. 139782

Rao, S.S.P., Huntley, M.H., Durand, N.C., Stamenova, E.K., Bochkov, I.D., Robinson, J.T., Sanborn, A.L., Machol, I., Omer, A.D., Lander, E.S., et al. (2014). A 3D map of the human genome at kilobase resolution reveals principles of chromatin looping. Cell 159, 1665–1680.

Rodman, T.C., Pruslin, F.H., Hoffmann, H.P., and Allfrey, V.G. (1981). Turnover of basic chromosomal proteins in fertilized eggs: a cytoimmunochemical study of events in vivo. J. Cell Biol. 90, 351–361.

Sanborn, A.L., Rao, S.S.P., Huang, S.-C., Durand, N.C., Huntley, M.H., Jewett, A.I., Bochkov, I.D., Chinnappan, D., Cutkosky, A., Li, J., et al. (2015). Chromatin extrusion explains key features of loop and domain formation in wild-type and engineered genomes. Proc. Natl. Acad. Sci. 112, 201518552.

Schwarzer, W., Abdennur, N., Goloborodko, A., Pekowska, A., Fudenberg, G., Loe-Mie, Y., Fonseca, N.A., Huber, W., Haering, C., Mirny, L., et al. (2016). Two independent modes of chromosome organization are revealed by cohesin removal. bioRxiv 94185.

Seitan, V.C., Faure, A.J., Zhan, Y., McCord, R.P., Lajoie, B.R., Ing-Simmons, E., Lenhard, B., Giorgetti, L., Heard, E., Fisher, A.G., et al. (2013). Cohesin-based chromatin interactions enable regulated gene expression within preexisting architectural compartments. Genome Res. 23, 2066–2077.

Sofueva, S., Yaffe, E., Chan, W.-C., Georgopoulou, D., Vietri Rudan, M., Mira-Bontenbal, H., Pollard, S.M., Schroth, G.P., Tanay, A., and Hadjur, S. (2013). Cohesin-mediated interactions organize chromosomal domain architecture. EMBO J. 32, 3119–3129.

Strom, A.R., Emelyanov, A. V., Mir, M., Fyodorov, D. V., Darzacq, X., and Karpen, G.H. (2017). Phase separation drives heterochromatin domain formation. Nature 547, 241–245.

Tachibana-Konwalski, K., Godwin, J., van der Weyden, L., Champion, L., Kudo, N.R., Adams, D.J., and Nasmyth, K. (2010). Rec8-containing cohesin maintains bivalents without turnover during the growing phase of mouse oocytes. Genes Dev. 24, 2505–2516.

Tardat, M., Albert, M., Kunzmann, R., Liu, Z., Kaustov, L., Thierry, R., Duan, S., Brykczynska, U., Arrowsmith, C.H., and Peters, A.H.F.M. (2015). Cbx2 Targets PRC1 to Constitutive Heterochromatin in Mouse Zygotes in a Parent-of-Origin-Dependent Manner. Mol. Cell 58, 157–171.

Tedeschi, A., Wutz, G., Huet, S., Jaritz, M., Wuensche, A., Schirghuber, E., Davidson, I.F., Tang, W., Cisneros, D.A., Bhaskara, V., et al. (2013). Wapl is an essential regulator of chromatin structure and chromosome segregation. Nature 501, 564–568.

Terekawa, T., Bisht, S., Eeftens, J., Dekker, C., Haering, C., and Greene, E. (2017). The Condensin Complex Is A Mechanochemical Motor That Translocates Along DNA. bioRxiv. 137711

Tjong, H., Gong, K., Chen, L., and Alber, F. (2012). Physical tethering and volume exclusion determine higher-order genome organization in budding yeast. Genome Res. 22, 1295–1305.

Tonkin, E.T., Wang, T.-J., Lisgo, S., Bamshad, M.J., and Strachan, T. (2004). NIPBL, encoding a homolog of fungal Scc2-type sister chromatid cohesion proteins and fly Nipped-B, is mutated in Cornelia de Lange syndrome. Nat. Genet. 36, 636–641.

Torres-Padilla, M.-E., Bannister, A.J., Hurd, P.J., Kouzarides, T., and Zernicka-Goetz, M. (2006). Dynamic distribution of the replacement histone variant H3.3 in the mouse oocyte and preimplantation embryos. Int. J. Dev. Biol. 50, 455–461.

Tran, N.T., Laub, M.T., and Le, T.B.K. (2017). SMC progressively aligns chromosomal arms in Caulobacter crescentus but is antagonized by convergent transcription. bioRxiv. 125344

Ulianov, S. V., Tachibana-Konwalski, K., and Razin, S. V. (2017). Single-cell Hi-C bridges microscopy and genome-wide sequencing approaches to study 3D chromatin organization. BioEssays 1700104.

Vietri Rudan, M., Barrington, C., Henderson, S., Ernst, C., Odom, D.T., Tanay, A., and Hadjur, S. (2015). Comparative Hi-C Reveals that CTCF Underlies Evolution of Chromosomal Domain Architecture. Cell Rep. 10, 1297–1309.

Wang, X., Brandão, H.B., Le, T.B.K., Laub, M.T., and Rudner, D.Z. (2017). Bacillus subtilis SMC complexes juxtapose chromosome arms as they travel from origin to terminus. Science. 355, 524–527.

Wendt, K.S., Yoshida, K., Itoh, T., Bando, M., Koch, B., Schirghuber, E., Tsutsumi, S., Nagae, G., Ishihara, K., Mishiro, T., et al. (2008). Cohesin mediates transcriptional insulation by CCCTC-binding factor. Nature 451, 796–801.

Zuin, J., Dixon, J.R., van der Reijden, M.I.J. a, Ye, Z., Kolovos, P., Brouwer, R.W.W., van de Corput, M.P.C., van de Werken, H.J.G., Knoch, T. a, van Ijcken, W.F.J., et al. (2013). Cohesin and CTCF differentially affect chromatin architecture and gene expression in human cells. Proc. Natl. Acad. Sci. U. S. A. 111, 996–1001.

